# Discovery and Quantification of Long-Range RNA Base Pairs in Coronavirus Genomes with SEARCH-MaP and SEISMIC-RNA

**DOI:** 10.1101/2024.04.29.591762

**Authors:** Matthew F. Allan, Justin Aruda, Jesse S. Plung, Scott L. Grote, Yves J. Martin des Taillades, Albéric A. de Lajarte, Mark Bathe, Silvi Rouskin

## Abstract

RNA molecules perform a diversity of essential functions for which their linear sequences must fold into higher-order structures. Techniques including crystallography and cryogenic electron microscopy have revealed 3D structures of ribosomal, transfer, and other well-structured RNAs; while chemical probing with sequencing facilitates secondary structure modeling of any RNAs of interest, even within cells. Ongoing efforts continue increasing the accuracy, resolution, and ability to distinguish coexisting alternative structures. However, no method can discover and quantify alternative structures with base pairs spanning arbitrarily long distances – an obstacle for studying viral, messenger, and long noncoding RNAs, which may form long-range base pairs.

Here, we introduce the method of Structure Ensemble Ablation by Reverse Complement Hybridization with Mutational Profiling (SEARCH-MaP) and software for Structure Ensemble Inference by Sequencing, Mutation Identification, and Clustering of RNA (SEISMIC-RNA). We use SEARCH-MaP and SEISMIC-RNA to discover that the frameshift stimulating element of SARS coronavirus 2 base-pairs with another element 1 kilobase downstream in nearly half of RNA molecules, and that this structure competes with a pseudoknot that stimulates ribosomal frameshifting. Moreover, we identify long-range base pairs involving the frameshift stimulating element in other coronaviruses including SARS coronavirus 1 and transmissible gastroenteritis virus, and model the full genomic secondary structure of the latter. These findings suggest that long-range base pairs are common in coronaviruses and may regulate ribosomal frameshifting, which is essential for viral RNA synthesis. We anticipate that SEARCH-MaP will enable solving many RNA structure ensembles that have eluded characterization, thereby enhancing our general understanding of RNA structures and their functions. SEISMIC-RNA, software for analyzing mutational profiling data at any scale, could power future studies on RNA structure and is available on GitHub and the Python Package Index.

## Introduction

Across all domains of life, RNA molecules perform myriad functions in development [1], immunity [2], translation [3], sensing [4, 5], epigenetics [6], cancer [7], and more. RNA also constitutes the genomes of many threatening viruses [8], including influenza viruses [9] and coronaviruses [10]. The capabilities of an RNA molecule depend not only on its sequence (primary structure) but also on its base pairs (secondary structure) and three-dimensional shape (tertiary structure) [11].

Although high-quality tertiary structures provide the most information, resolving them often proves difficult or impossible with mainstay methods used for proteins [12]. Consequently, the world’s largest database of tertiary structures – the Protein Data Bank [13] – has accumulated only 1,839 structures of RNAs (compared to 198,506 of proteins) as of February 2024. Worse, most of those RNAs are short: only 119 are longer than 200 nt; of those, only 24 are not ribosomal RNAs or group I/II introns. Due partly to the paucity of non-redundant long RNA structures, methods of predicting tertiary structures for RNAs lag far behind those for proteins [14].

The situation is only marginally better for RNA secondary structures. If a diverse set of homologous RNA sequences is available, a consensus secondary structure can often be predicted using comparative sequence analysis, which has accurately modeled ribosomal and transfer RNAs, among others [15]. A formalization known as the covariance model [16] underlies the widely-used Rfam database [17] of consensus secondary structures for 4,170 RNA families (as of version 14.10). Although extensive, Rfam contains no protein-coding sequences (with some exceptions such as frameshift stimulating elements) and provides only one secondary structure for each family, even though many RNAs fold into multiple functional structures [18, 19]). Each family also models only a short segment of a full RNA sequence; for coronaviruses, existing families encompass the 5’ and 3’ untranslated regions, the frameshift stimulating element, and the packaging signal, which collectively constitute only 3% of the genomic RNA.

Predicting secondary structures faces two major obstacles due to the scarcity of high-quality RNA structures, particularly for RNAs longer than 200 nt (including long non-coding [20], messenger [21], and viral genomic [22] RNAs). First, prediction methods trained on known RNA structures are limited to small, low-diversity training datasets (generally of short sequences), which causes overfitting and hence inaccurate predictions for dissimilar RNAs (including longer sequences) [23, 24]. Second, without known secondary structures of many diverse RNAs, the accuracy of any prediction method cannot be properly bench-marked [21, 25]. For these reasons, and because thermodynamic-based models also tend to be less accurate for longer RNAs [22] and base pairs spanning longer distances [26], predicting secondary structures of long RNAs remains unreliable.

The most promising methods for determining the structures of long RNAs use experimental data. Chemical probing experiments involve treating RNA with reagents that modify nucleotides depending on the local secondary structure; for instance, dimethyl sulfate (DMS) methylates adenosine (A) and cytidine (C) residues only if they are not base-paired [27]. Modern methods use reverse transcription to encode modifications of the RNA as mutations in the cDNA, followed by next-generation sequencing – a strategy known as mutational profiling (MaP) [28, 29]. A key advantage of MaP is that the sequencing reads can be clustered to detect multiple secondary structures in an ensemble [30, 31]. Determining the base pairs in those structures still requires structure prediction [32], although incorporating chemical probing data does improve accuracy [33, 34].

Several experimental methods have been developed to find base pairs directly, with minimal reliance on structure prediction. M2-seq [35] introduces random mutations before chemical probing to detect correlated mutations between pairs of bases, which indicates the bases interact. However, alternative structures complicate the data analysis [36], and detectible base pairs can be no longer than the sequencing reads (typically 300 nt). For long-range base pairs, many methods involving crosslinking, proximity ligation, and sequencing have been developed [37]. These methods can find base pairs spanning arbitrarily long distances – as well as between different RNA molecules – but cannot resolve single base pairs or alternative structures. Detecting, resolving, and quantifying alternative structures with base pairs that span arbitrarily long distances remains an open challenge.

Here, we introduce “Structure Ensemble Ablation by Reverse Complement Hybridization with Mutational Profiling” (SEARCH-MaP), an experimental method to discover RNA base pairs spanning arbitrarily long distances. We also develop the software “Structure Ensemble Inference by Sequencing, Mutation Identification, and Clustering of RNA” (SEISMIC-RNA) to analyze MaP data and resolve alternative structures. Using SEARCH-MaP and SEISMIC-RNA, we discover an RNA structure in severe acute respiratory syndrome coronavirus 2 (SARS-CoV-2) that comprises dozens of long-range base pairs and folds in nearly half of genomic RNA molecules. We show that it inhibits the folding of a pseudoknot that stimulates ribosomal frameshifting [38, 39], hinting a role in regulating viral protein synthesis. We find similar structures in other SARS-related viruses and transmissible gastroenteritis virus (TGEV), suggesting that long-range base pairs involving the frameshift stimulation element are a general feature of coronaviruses. In addition to revealing new structures in coronaviral genomes, our findings show how SEARCH-MaP and SEISMIC-RNA can resolve secondary structure ensembles of long RNA molecules – a necessary step towards a true “AlphaFold for RNA” [14].

## Results

### SEISMIC-RNA analyzes chemical probing data and resolves alternative RNA structures

SEISMIC-RNA is all-in-one software for processing and graphing mutational profiling data (from e.g. DMS-MaPseq [29] and SHAPE-MaP [28] experiments), inferring alternative structures, and modeling secondary structures (Figure 1a). Designed for modern high-throughput experiments, SEISMIC-RNA can process multiple samples in parallel, quickly, with low memory and storage footprints. SEISMIC-RNA ensures data quality with new algorithms that detect ambiguous base calls (e.g. deletions within repetitive sequences) and mask unusable reads and positions (e.g. with insufficient coverage). To identify alternative structures, SEISMIC-RNA introduces a new clustering algorithm – similar to DREEM [30] yet able to cluster RNAs longer than the reads themselves – enabling analyses of structure ensembles in long transcripts. On simulated datasets, SEISMIC-RNA detected the correct number of alternative structures given sufficient reads (Figure 1b, c) and inferred their proportions and chemical reactivities accurately (Supplementary Figure 1). SEISMIC-RNA can use the chemical reactivities to model secondary structures and generate a variety of graphs – including correlations between samples and receiver operating characteristic (ROC) curves. The entire workflow can be run on the command line with one command, for ease of use; while a second Python interface enables highly customized analyses.

**Figure 1:**
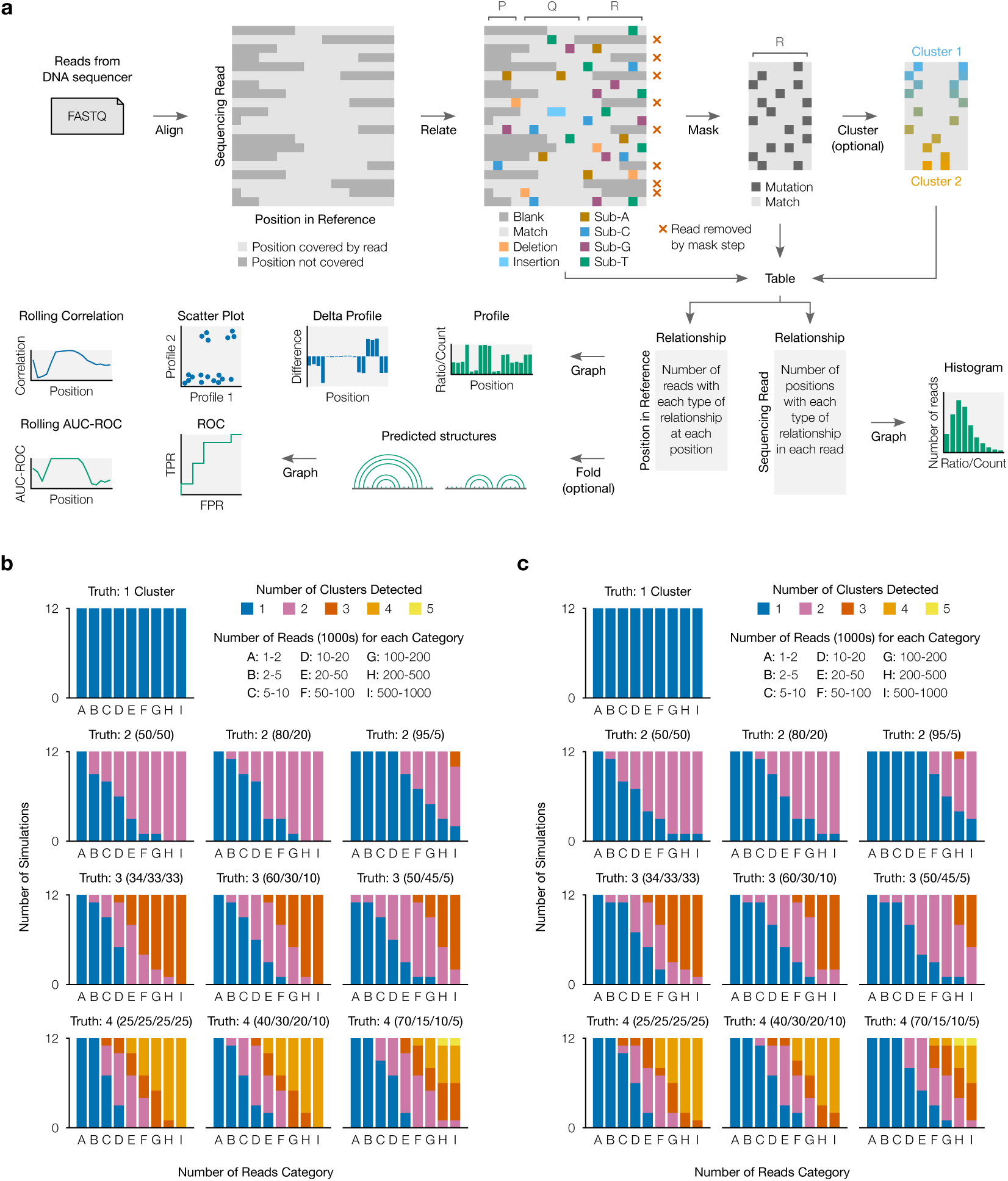
SEISMIC-RNA software. **(a)** SEISMIC-RNA first trims sequencing reads with Cutadapt [40] and aligns them to reference sequence(s) with Bowtie 2 [41]). For every read, the relationship (i.e. match, substitution, deletion, etc.) to each position in the reference is determined. Relationships are classified as mutated, matched, or uninformative; and positions and reads are masked by user-specified filters. Reads can then be clustered to reveal alternative structures. The types of relationships at each position and in each read are tabulated. SEISMIC-RNA can use these tables to predict RNA secondary structures via RNAstructure [42, 33] or graph mutational profiles, scatter plots, receiver operating characteristic (ROC) curves, and more. **(b)** To validate the clustering algorithm, datasets were simulated with one to four “true” alternative structure(s) in three mixing proportions and with 1,000 to 1,000,000 full-length reads. For each number of reads, 12 unique 280-nt RNAs were simulated and processed with SEISMIC-RNA. The number of clusters (up to 5) detected by each simulation is shown. **(c)** Same as (b), but reads were simulated with random 5’ and 3’ ends rather than fully covering the RNA.

### SEARCH-MaP detects long-range base pairs in RNA structure ensembles

In a SEARCH-MaP experiment, an RNA is assumed to fold into an ensemble of one or more structures (Figure 2a). Searching for base pairs involving part of the RNA – in this example, section P – begins by binding an antisense oligonucleotide (ASO) to that part, which prevents it from base pairing (Figure 2b). Separately, the RNA is chemically probed with (+ASO) and without (no-ASO) the ASO, followed by mutational profiling (MaP) and sequencing (e.g. DMS-MaPseq [29]).

**Figure 2:**
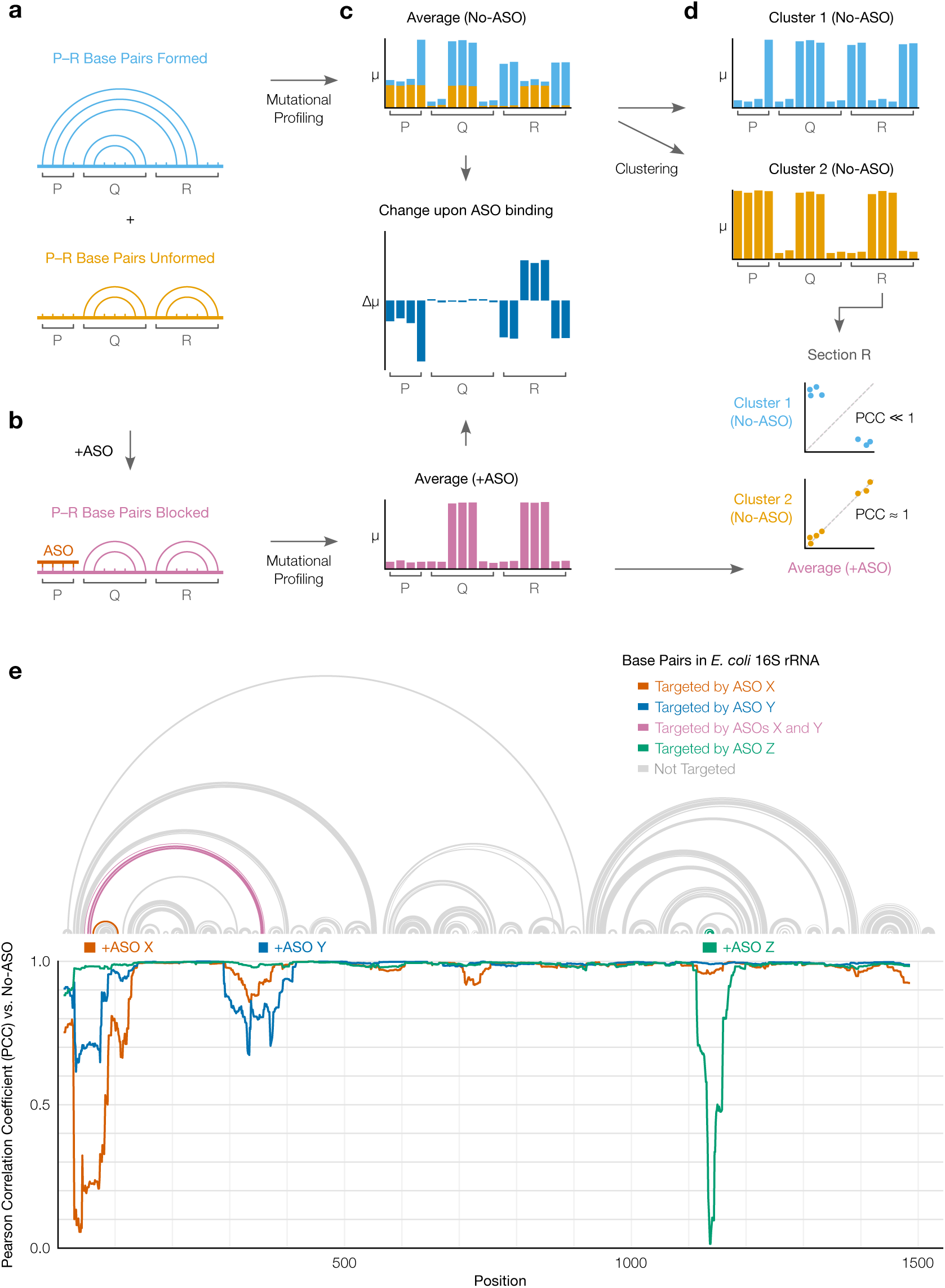
SEARCH-MaP strategy and validation. **(a)** This RNA comprises three sections (P, Q, and R) and folds into an ensemble of two structures: one in which base pairs between P and R form and one in which they do not. **(b)** Hybridizing an ASO to P blocks it from base-pairing with R. **(c)** Mutational profiles with (+ASO) and without (no-ASO) the ASO, computed as ensemble averages with SEISMIC-RNA. The *x*-axis is the position in the RNA sequence; the *y*-axis is the fraction of mutated bases (*µ*) at the position. Each bar in the no-ASO profile is drawn in two colors merely to illustrate how many mutations at each position come from each structure; in a real experiment, this information would not exist before clustering. The change upon ASO binding indicates the difference in the fraction of mutated bases (Δ*µ*) between the +ASO and no-ASO conditions. **(d)** Mutational profiles of two clusters (top) obtained by clustering the no-ASO ensemble in (c) using SEISMIC-RNA, and scatter plots comparing the mutational profiles (bottom) between the +ASO ensemble average (*x*-axis) and each cluster (*y*-axis); each point represents one base in section R. The expected Pearson correlation coefficient (PCC) is shown beside each scatter plot. **(e)** Validation of SEARCH-MaP on 16S ribosomal RNA (rRNA) from *E. coli*. The canonical secondary structure [15] is shown above the target location of each ASO. The rolling (window = 45 nt) PCC of the DMS reactivities with each ASO versus without any ASOs is graphed below.

Each structure theoretically has its own mutational profile, which is not directly observable because all structures are physically mixed in an experiment [43]. The only directly observable mutational profile is of the “ensemble average” – the average of the structures’ (unobserved) mutational profiles, weighted by the their (unobserved) proportions (Figure 2c, top). The structure – and thus mutational profile – of section R changes when it base-pairs with P; by preventing such base-pairing, the ASO changes the ensemble average of R (Figure 2c, middle). However, the ASO has no effect on section Q because Q never base-pairs with P. Therefore, one can deduce that P base-pairs with R (but not with Q) because hybridizing an ASO to P alters the mutational profile of R (but not of Q).

Furthermore, the mutational profile of the structure in which P and R base-pair can be determined, even without knowing the exact base pairs. This step involves clustering the no-ASO ensemble into two mutational profiles over section R – each corresponding to one structure – and comparing them to the +ASO ensemble average (Figure 2d). Because the ASO blocks the P–R base pairs, the +ASO mutational profile will correlate better with that of the structure where P and R do not base-pair; in this case, cluster 2 correlates better. Therefore, the mutational profile of cluster 1 corresponds to the structure where P and R base-pair.

We confirmed that SEARCH-MaP can detect known long-range base pairs in 16S ribosomal RNA from *E. coli* (Figure 2e). Two ASOs, X and Y, targeted opposite sides of a known stem spanning just over 300 nt. Adding ASO X caused the correlation versus the no-ASO sample to dip below 0.1 at the target site as well as below 0.9 on the other side of the stem, confirming that we could detect this structure. Adding ASO Y likewise dropped the correlation at both its target site and the other side of the stem to below 0.7; no other sites were perturbed. As another control, ASO Z – which targeted the tip of a stem loop rather than one side – perturbed the correlation only at the target site, as expected. These results show that the SEARCH-MaP strategy of adding ASOs followed by mutational profiling can indeed detect known long-range base pairs.

### SEARCH-MaP detects and quantifies long-range base pairing in SARS-CoV-2

Aside from ribosomes, many of the best-characterized functional long-range RNA base pairs occur in the genomes of RNA viruses [44]. In coronaviruses, the first open reading frame (ORF1) contains a frameshift stimulation element (FSE) that makes a fraction of ribosomes slip into the −1 reading frame, bypass a 0-frame stop codon, and translate to the end of ORF1 [45]. Every coronaviral FSE contains a “slippery site” (UUUAAAC) and a structure characterized as a pseudoknot in multiple species [46, 47, 48]. For SARS coronavirus 2 (SARS-CoV-2), 80-90 nt segments of the core FSE have been shown to fold into a pseudoknot with three stems [39, 49, 50]. However, in intact SARS-CoV-2, the FSE adopts a different structure: one whose DMS reactivities could be recapitulated *in vitro* with a 2,924 nt segment but not a 283 nt segment, suggesting long-range base pairs [51]. To search for long-range base pairs explaining these results, we performed SEARCH-MaP on the 2,924 nt segment after adding each of thirteen groups of DNA ASOs tiling the entire transcript (Figure 3a). We used one pair of RT-PCR primers flanking each ASO target site to confirm binding (Supplementary Figure 2) and another pair flanking the 283 nt section of the FSE to assess its structure (Supplementary Figure 3).

**Figure 3:**
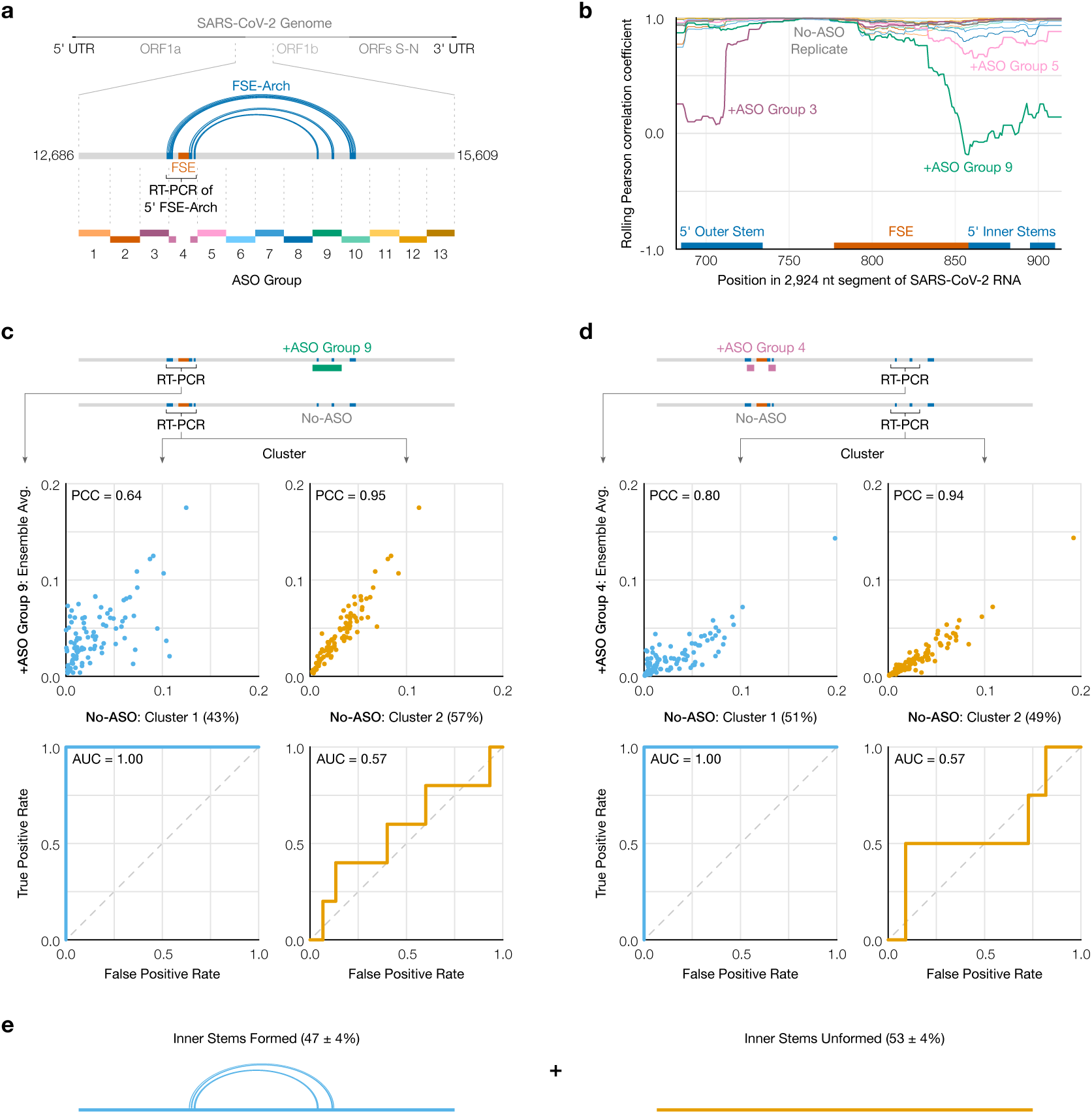
SEARCH-MaP of long-range base pairs involving the SARS-CoV-2 FSE. **(a)** The 2,924 nt segment of the SARS-CoV-2 genome containing the frameshift stimulation element (FSE) and model of the FSE-arch [52]. The target site for each group of antisense oligonucleotides (ASOs) is indicated by dotted lines; lengths are to scale. **(b)** Rolling (window = 45 nt) Pearson correlation coefficient (PCC) of DMS reactivities over the 5’ FSE-arch between each +ASO sample and a no-ASO control. Each curve represents one ASO group, colored as in (a); groups 4 and 13 are not shown. Locations of the FSE and the outer and inner stems of the 5’ FSE-arch are also indicated. **(c)** (Top) Scatter plots of DMS reactivities over the 5’ FSE-arch comparing each cluster of the no-ASO sample to the sample with ASO group 9, with PCC indicated; each point is one base in the 5’ FSE-arch. (Bottom) Receiver operating characteristic (ROC) curves comparing each cluster of the no-ASO sample to the two inner stems of the FSE-arch, with area under the curve (AUC) indicated. **(d)** Like (c) but over the 3’ FSE-arch, and comparing to the sample with ASO group 4. One highly reactive outlier was excluded when calculating PCC (which is sensitive to outliers) but included in the ROC (which is robust). **(e)** Model of the inner two stems in the ensemble of structures formed by the 2,924 nt segment.

To quantify structural changes over the FSE, we calculated the rolling Pearson correlation coefficient (PCC) of the DMS reactivities between each sample and a no-ASO control (Figure 3b). The rolling PCC of a no-ASO replicate remained between 0.93 and 1.00 (mean = 0.97), confirming the DMS reactivities were reproducible. Besides ASO group 4 (targeting the FSE itself), the FSE structure was most perturbed by ASO group 9, targeting about 1 kb downstream. Intervening ASO groups 6, 7, and 8 had no effect on the FSE.

This result is consistent with a proposed structure named the “FSE-arch” [52] comprising three long-range stems (Figure 3a). The two inner stems connect the FSE (on the 5’ side) to the target site of ASO group 9 (on the 3’ side); thus, ASO group 9 would have disrupted them and perturbed the FSE, as we observed. For the outer stem, the 3’ side lies within the target site for ASO group 10, but this group had no effect: the outer stem apparently did not form. These results suggest both inner stems (but not the outer stem) of the proposed FSE-arch [52] exist, explaining our previous finding that a 283 nt segment was insufficient to recapitulate the FSE structure *in vitro* [51].

To determine in what fraction of molecules the two inner stems of the FSE-arch form, we clustered reads from the 5’ side of the FSE-arch for the no-ASO control. We found two clusters with a 43/57% split and – to determine if they corresponded to the two inner stems formed versus unformed – compared their DMS reactivities to those after adding ASO group 9, which would block the two inner stems (Figure 3c, top). cluster 2 had similar DMS reactivities (PCC = 0.95), indicating it corresponds to the stems unformed. Meanwhile, the DMS reactivities of cluster 1 differed (PCC = 0.64), suggesting it corresponds to the stems formed.

To further support this result, we leveraged the preexisting model of the FSE-arch [52]. If cluster 1 did correspond to the two inner stems formed, its DMS reactivities would agree well with their structures (i.e. paired and unpaired bases should have low and high reactivities, respectively), while those of cluster 2 would agree less. We quantified this agreement using receiver operating characteristic (ROC) curves (Figure 3c, bottom). The area under the curve (AUC) for cluster 1 was 1.00, indicating perfect agreement with the two inner stems of the FSE-arch; while that of cluster 2 was 0.57, close to no agreement (0.50). This result further supports that clusters 1 and 2 correspond to the two inner stems formed and unformed, respectively.

If the RNA did exist as an ensemble of the two inner stems formed and unformed, the 3’ side of the FSE-arch would also cluster into states with the two inner stems formed and unformed. We thus RT-PCRed and clustered the 3’ FSE-arch in the no-ASO control and +ASO group 4 and found – similar to the previous result – that the DMS reactivities after blocking the 5’ FSE-arch with ASO group 4 resembled those of cluster 2 (PCC = 0.94) but not cluster 1 (PCC = 0.80), while the structure of the two inner stems agreed with cluster 1 (AUC = 1.00) but not cluster 2 (AUC = 0.57) (Figure 3d). We concluded that the RNA exists as an ensemble of structures in which the two inner stems of the FSE-arch form in 47% ± 4% of molecules (Figure 3e).

### The long-range stems compete with the frameshift pseudoknot in SARS-CoV-2

To determine if the FSE forms other long-range stems, in lieu of the original outer stem of the FSE-arch [52], we modeled a 1,799 nt segment centered on the FSE-arch. Although computationally predicting long-range base pairs is notoriously unreliable [26, 22], we speculated that we could improve accuracy by incorporating the DMS reactivities of cluster 1 on both sides of the FSE-arch (Supplementary Figure 4). For the innermost stem – which we call long stem 1 (LS1) – nine of thirteen structures (69%) predicted using the cluster 1 DMS reactivities contained LS1, compared to five of eleven (45%) using the ensemble average and four of twenty (20%) using no DMS reactivities. For the second-most inner stem (LS2), eight structures (62%) predicted using cluster 1 contained LS2, while none did using average or no DMS reactivities. Thus, the DMS reactivities corresponding to the long-range cluster enabled predicting the long-range stems more consistently, allowing us to refine our model of the long-range stems.

Our refined model based on the long-range cluster (Figure 4a) included not only the two inner stems of the FSE-arch – LS1 and LS2a/b – but also two long stems (LS3a/b and LS4) absent from the original FSE-arch model [52], as well as alternative stem 1 (AS1) [51]. To verify this model, we performed SEARCH-MaP on the 1,799 nt segment using 15-20 nt LNA/DNA mixmer ASOs for single-stem precision (Supplementary Table 3). Each ASO targeted the 3’ side of one stem, and we measured the change in DMS reactivities of the FSE (Figure 4b). ASOs targeting the 3’ sides of LS1 and LS2a perturbed the DMS reactivities in exactly the expected locations on the 5’ sides. An ASO for the 3’ side of LS2b perturbed the FSE with more off-target effects, likely because this stem overlaps with pseudoknot stem 2 (PS2). Blocking LS3b also resulted in a main effect around the intended location, with one off-target effect upstream, suggesting that other base pairs between the pseudoknot and this upstream region may exist. Therefore, stems LS1, LS2a/b, and LS3b do exist – at least in a portion of the ensemble.

**Figure 4:**
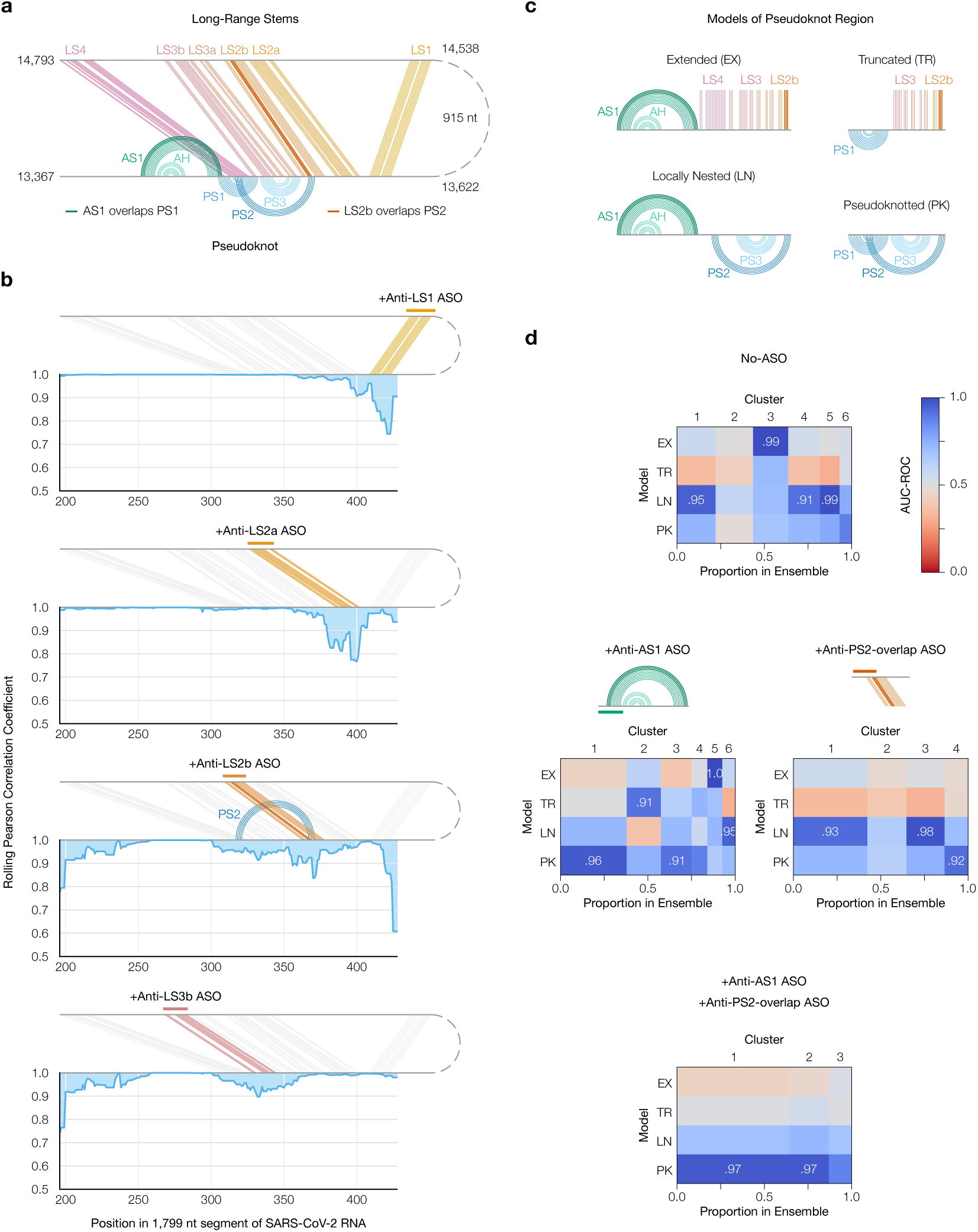
Refinement of the long-range structure model and competition with the frameshift pseudoknot. **(a)** Refined model of the long-range stems (minimum free energy prediction based on cluster 1 in Figure 3c and d) including alternative stem 1 (AS1) [51]; the attenuator hairpin (AH) [53]; and long stems LS1, LS2a/b, LS3a/b, and LS4. Locations of pseudoknot stems PS1, PS2, and PS3 [39] are also shown; as are the base pairs they overlap in AS1 and LS2b. **(b)** Rolling (window = 21 nt) Pearson correlation coefficient of DMS reactivities between each +ASO sample and a no-ASO control; base pairs targeted by each ASO are colored. **(c)** Models of possible structures for the FSE, by combining non-overlapping stems from (a). **(d)** Heatmaps comparing models in (c) to clusters of DMS reactivities over positions 305-371 via the area under the receiver operating characteristic curve (AUC-ROC). AUC-ROCs at least 0.90 are annotated. Cluster widths indicate proportions in the ensemble.

LS2b, LS3, and LS4 of the refined model overlap all three stems of the pseudoknot (PS1, PS2, and PS3) that stimulates frameshifting [39]. To test whether these long stems actually compete with the pseudoknot, we made four possible models of the FSE structure by combining mutually compatible stems (Figure 4c). Then, we clustered the 1,799 nt segment without ASOs up to 6 clusters – the maximum number reproducible between replicates (Supplementary Figure 5a) – and compared each cluster to each structure model using the area under the receiver operating characteristic curve (AUC-ROC) over positions 305-371, spanned by the pseudoknot (Figure 4d, top). We considered a cluster and model to be “consistent” if the AUC-ROC was at least 0.90. The locally nested model (AS1 plus PS2 and PS3) was consistent with three clusters totaling 52% of the ensemble, while the extended model (AS1 plus all long-range stems) was consistent with one cluster (20%). No clusters were fully consistent with the pseudoknotted model, though the least-abundant cluster (7%) came close with an AUC-ROC of 0.88. The remaining cluster (21%) was not consistent with any model, suggesting that the ensemble contains structures beyond those in Figure 4c.

Adding an ASO targeting the 5’ side of AS1 reduced the proportion of AS1-containing states (extended and locally nested) from 72% to 16% (Figure 4d, left; Supplementary Figure 5b). In their place emerged clusters consistent with the pseudoknotted and truncated models, constituting 56% and 20% of the ensemble, respectively. Meanwhile, adding an ASO that blocked the part of LS2b that overlaps PS2 eliminated the extended state (which includes LS2b) and produced one cluster (13%) consistent with the pseudoknotted model (Figure 4d, right; Supplementary Figure 5c). Adding both ASOs simultaneously collapsed the ensemble into three clusters of which two (87%) were highly consistent with the pseudo-knotted model (Figure 4d, bottom; Supplementary Figure 5d). Since blocking the PS2-overlapping portion of LS2b increased the proportion of clusters consistent (or nearly so) with the pseudoknotted model – both alone and combined with the anti-AS1 ASO – the long-range stems did appear to outcompete the pseudoknot.

**Figure 5:**
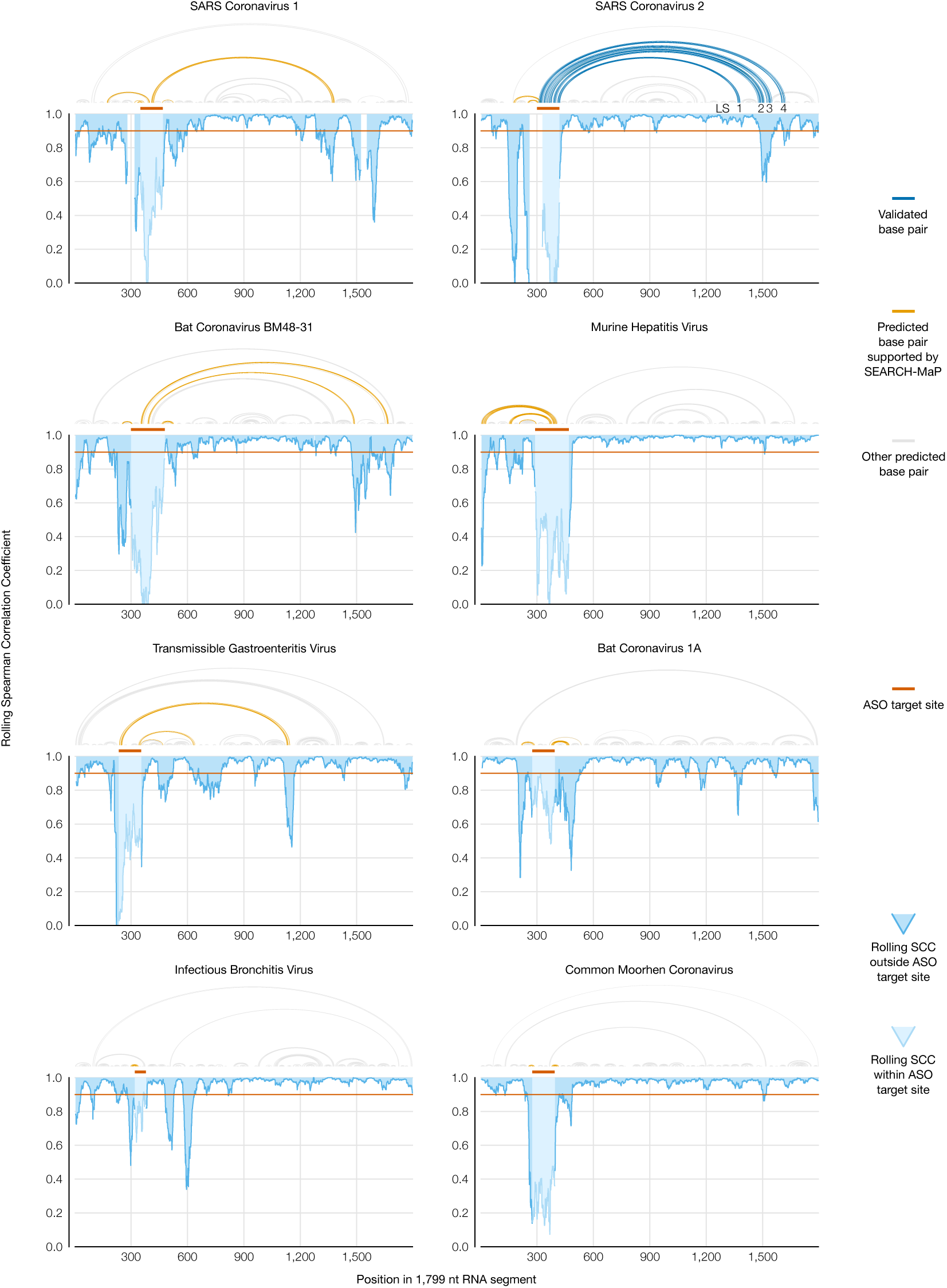
Evidence for long-range RNA–RNA base pairs involving the FSE in four additional coronaviruses. Rolling (window = 45 nt) Spearman correlation coefficient (SCC) of DMS reactivities between the +ASO and no-ASO samples for each 1,799 nt segment of a coronaviral genome. The target site of each ASO is highlighted on the SCC data and shown above each graph. Structures predicted with RNAstructure [42] using no-ASO ensemble average DMS reactivities as constraints [33] are drawn above each graph; base pairs connecting the ASO target site to an off-target position with SCC less than 0.9 are colored. For SARS-CoV-2, the refined model (Figure 4a) is also drawn, with LS1-LS4 labeled.

### Frameshift stimulating elements of multiple coronaviruses form long-range base pairs

We hypothesized that other coronaviruses would also feature long-range base pairs involving their FSEs. To search for such coronaviruses computationally, we predicted structures of 2,000 nt sections surrounding the FSEs of all 62 complete coronavirus genomes in the NCBI Reference Sequence Database [54] (Supplementary Figure 6). We selected ten coronaviruses, at least one from each genus (Supplementary Figure 7a), based on with which bases the FSE was predicted to base-pair. In *Betacoronavirus*, SARS coronaviruses 1 (`NC_004718.3`) and 2 (`NC_045512.2`) and bat coronavirus BM48-31 (`NC_014470.1`) because they clustered into their own structural outgroup; MERS coronavirus (`NC_019843.3`), predicted to pair with positions 510-530; and human coronavirus OC43 (`NC_006213.1`) and murine hepatitis virus strain A59 (`NC_048217.1`), both predicted to pair with positions 10-20. In *Alphacoronavirus*, transmissible gastroenteritis virus (`NC_038861.1`) and bat coronavirus 1A (`NC_010437.1`), predicted to pair with positions 440-460 and 350-360, respectively. In *Gammacoronavirus*, avian infectious bronchitis virus strain Beaudette (`NC_001451.1`), predicted to pair with positions 330-350. And in *Deltacoronavirus*, common moorhen coronavirus HKU21 (`NC_016996.1`), which had the most promising long-range base pairs in this genus.

We screened each of these ten coronaviruses for long-range base pairs with the FSE by comparing the DMS reactivities of a 239 nt segment comprising the FSE with minimal flanking sequences and a 1,799 nt segment encompassing the FSE and all sites with which it was predicted to interact. All coronaviruses except MERS coronavirus and human coronavirus OC43 showed differences in their DMS reactivities between the 239 and 1,799 nt segments (Supplementary Figure 7b), suggesting their FSEs formed long-range base pairs.

To locate base pairs involving the FSE in each coronavirus, we performed SEARCH-MaP on the 1,799 nt RNA segment using DNA ASOs targeting the vicinity of the FSE (Figure 5). The rolling Spearman correlation coefficient (SCC) between the +ASO and no-ASO mutational profiles dipped below 0.9 at the ASO target site in every coronavirus segment, confirming the ASOs bound and altered the structure. To confirm we could detect long-range base pairs, we compared the rolling SCC for the SARS-CoV-2 segment to our refined model of the FSE structure (Figure 5, blue). The SCC dipped below 0.9 at positions 1,483-1,560 and at 1,611-1,642, coinciding with stems LS2-LS3 (positions 1,476-1,550 within the 1,799 nt segment) and stem LS4 (positions 1,600-1,622). These dips were the two largest downstream of the FSE; although others (corresponding to no known base pairs) existed, they were barely below 0.9 and could have resulted from base pairing between these and other (non-FSE) regions. Near LS1 (positions 1,367-1,381), the SCC dipped only slightly to 0.95, presumably because LS1 is the smallest (15 nt) and most isolated long-range stem. Therefore, this method was sensitive enough to detect all but the smallest long-range stem, and specific enough that the two largest dips corresponded to validated long-range stems.

We found similar long-range stems in SARS-CoV-1 and another SARS-related virus, bat coronavirus BM48-31. Both viruses showed dips in SCC at roughly the same positions as LS2-LS4 in SARS-CoV-2, indicating homologous structures. SARS-CoV-1 also had a wide dip below 0.9 at positions 1,284-1,394, corresponding to a homologous LS1. Thus, three SARS-related viruses share these long-range stems involving the FSE, hinting at a conserved function.

In every other species except common moorhen coronavirus, we found prominent dips in SCC at least 200 nt from the ASO target site. To determine what long-range base pairs could have caused those dips, we used RNAstructure Fold [42] guided by the DMS reactivities of the no-ASO ensemble average [33]. We obtained models consistent with the SEARCH-MaP data for both murine hepatitis virus and transmissible gastroenteritis virus (Figure 5, orange); although the algorithm did not successfully predict the long-range stems LS2-LS4 in SARS-CoV-1, likely due to the unreliability of long-range structure prediction [22] and our reliance on ensemble average (rather than clustered) DMS reactivities (Supplementary Figure 4). We conclude that long-range base pairing involving the FSE occurs more widely than in just SARS-CoV-2, including in the genus *Alphacoronavirus*.

### Structure of the full TGEV genome in ST cells supports long-range base pairing involving the FSE

Transmissible gastroenteritis virus (TGEV) is a strain of *Alphacoronavirus 1* [55] that infects pigs and causes vomiting and diarrhea, often fatally [56]. Due to the impacts of TGEV [56] and our evidence of long-range base pairs, we sought to model the genomic secondary structures of live TGEV. We began by infecting ST cells with TGEV and performing DMS-MaPseq [29] (Figure 6a). The DMS reactivities over the full genome were consistent between technical and biological replicates (PCC = 0.94-0.95, Supplementary Figure 8a and b), albeit not with the 1,799 nt segment *in vitro* (PCC = 0.76), which showed that verifying the long-range stem in live TGEV would be necessary (Supplementary Figure 8c).

**Figure 6:**
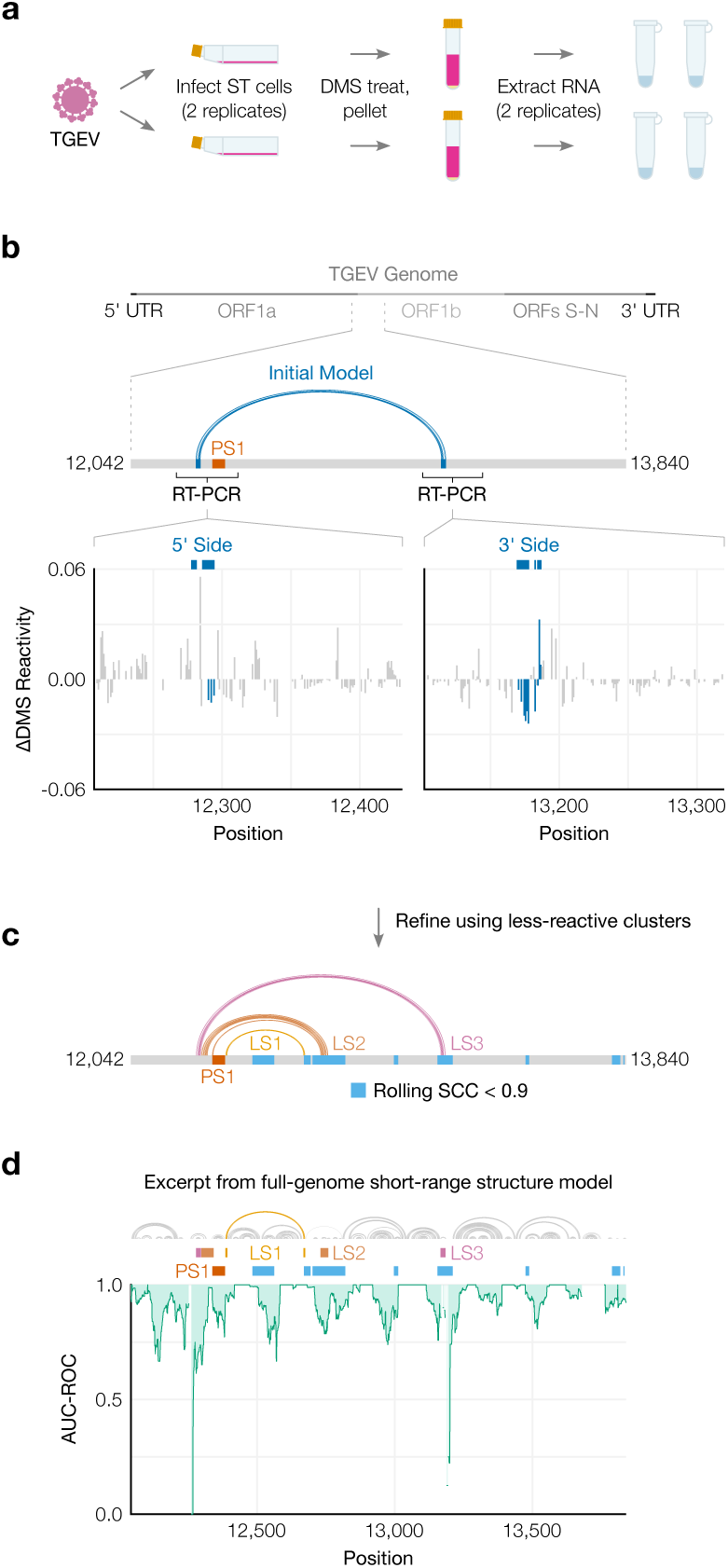
Genomic secondary structure of live TGEV. **(a)** Schematic of the experiment in which two biological replicates of ST cells were infected with TGEV, DMS-treated, and pelleted. Cell pellets were divided into two technical replicates prior to extraction of DMS-modified RNA. **(b)** Differences in DMS reactivities between the two clusters on each side of the long-range stem. Each bar represents one base. Bases are shaded dark blue if they pair in the initial model of the long-range stem (from Figure 5), shown above along with its location in the full genome. The locations of FSE pseudoknot stem 1 (PS1) and the regions amplified for clustering are also indicated. **(c)** Refined model of the long-range stem in TGEV based on the DMS reactivities of the less-reactive cluster from both sides. Long stems 1 (LS1), 2 (LS2), and 3 (LS3) are labeled. For comparison with the regions of the 1,799 nt segment perturbed by the ASO (Figure 5), positions after the FSE where the Spearman correlation coefficient (SCC) dipped below 0.9 are shaded light blue. **(d)** Rolling AUC-ROC (window = 45 nt) between the full-genome DMS reactivities and full-genome secondary structure modeled from the DMS reactivities (maximum 300 nt between paired bases). The structure model is drawn above the graph. Only positions 12,042-13,840 are shown here. For comparison, the locations of PS1, LS1, LS2, LS3, and dips in SCC after the FSE are also indicated.

To quantify the long-range base pairs, we RT-PCRed the 5’ and 3’ sides, confirmed their DMS reactivities were consistent with the full genome’s (Supplementary Figure 8d), and clustered both sides (Figure 6b). Bases involved in the predicted long-range pairs were generally less DMS-reactive in cluster 2 (Figure 6b, Supplementary Figure 9a), which we hypothesized corresponded to the long-range base pairs forming. In support, the long-range stem (hereafter, LS3) appeared in structures modeled from the DMS reactivities from cluster 2 on both sides (Figure 6c), but not cluster 1 (Supplementary Figure 9b). The refined model based on cluster 2 included another long-range stem, LS2, which was also supported by a dip in the rolling SCC (Figure 6c).

We used the ensemble average DMS reactivities to produce one “ensemble average” model of short-range (up to 300 nt) base pairs in the full TGEV genome (Supplementary Figure 10). To verify the model quality, we confirmed that the modeled structure of the first 330 nt included the highly conserved stem loops SL1, SL2, SL4, and SL5a/b/c in the 5’ UTR [10] (Supplementary Figure 11a) and was consistent with the DMS reactivities (AUC-ROC = 0.97) (Supplementary Figure 11b). The AUC-ROC was lower in many locations throughout the rest of the genome (Supplementary Figure 10), indicating that a single secondary structure consistent with the ensemble average DMS reactivities could not be found – which suggests alternative structures, long-range base pairs, or both [51]. Accordingly, we found a large dip in AUC-ROC just upstream of the FSE, centered on the 5’ ends of LS2 and LS3, as well as smaller dips at the 3’ ends of both stems (Figure 6d). In fact, at or near every location that SEARCH-MaP had evidenced to interact with the FSE – where the rolling SCC had dipped – the AUC-ROC also dipped. This finding supports that regions with low AUC-ROC in general are good starting points for investigating alternative structures and long-range base pairing with SEARCH-MaP.

## Discussion

In this work, we developed SEARCH-MaP and SEISMIC-RNA and applied them to detect structural ensembles involving long-range base pairs in SARS-CoV-2 and other coronaviruses. Previous studies have demonstrated that binding an ASO to one side of a long-range stem would perturb the chemical probing reactivities of the other side [57, 58, 59]. Here, we separated and identified the reactivities corresponding to long-range stems formed and unformed. This advance enables isolating the reactivities of the long-range stem formed – on not just one but both sides of the stem, linking corresponding alternative structures over distances much greater than the length of a read, which has not been possible in previous studies [30, 31]. Using the linked reactivities from both sides of a long-range stem, its secondary structure can be modeled more accurately than would be possible using the ensemble average reactivities, as we have done for SARS-CoV-2 (Supplementary Figure 4) and TGEV (Supplementary Figure 9).

SEISMIC-RNA builds upon our previous work, the DREEM algorithm [30]. Here, we have optimized the algorithm to run approximately 10-20 times faster and built an entirely new workflow around it for aligning reads, calling mutations, masking data, and outputting a variety of graphs. SEISMIC-RNA can process data from any mutational profiling experiment, including DMS-MaPseq [29] and SHAPE-MaP [28], not just SEARCH-MaP. The software is available from the Python Package Index (pypi.org/project/seismic-rna) and GitHub (github.com/rouskinlab/seismic-rna) and can be used as a command line executable program (`seismic`) or via its Python application programming interface (`import seismicrna`).

We envision SEARCH-MaP and SEISMIC-RNA bridging the gap between broad and detailed investigations of RNA structure. Other methods such as proximity ligation [60, 61, 62, 63, 64] provide broad, transcriptome-wide information on RNA structure and could be used as a starting point to find structures of interest for deeper investigation with SEARCH-MaP/SEISMIC-RNA. Indeed, the first evidence of the FSE-arch in SARS-CoV-2 came from such a study [52]. To investigate RNA structures in detail, M2-seq [35] and related methods [36] can pinpoint base pairs with up to single-nucleotide resolution and minimal need for structure prediction. However, base pairs are detectible only if the paired bases occur on the same sequencing read, which restricts their spans to at most the read length (typically 300 nt). Because the capabilities of M2-seq and SEARCH-MaP complement each other, they could be integrated: first SEARCH-MaP/SEISMIC-RNA to discover, quantify, and model long-range base pairs; then M2-seq for short-range base pairs. By providing the missing link – structure ensembles involving long-range base pairs – SEARCH-MaP and SEISMIC-RNA could combine broad and detailed views of RNA structure into one coherent model.

To understand structures of long RNA molecules, SEARCH-MaP and SEISMIC-RNA could also be used to validate predicted secondary structures and benchmark structure prediction algorithms. Algorithms that predict secondary structures achieve lower accuracies for longer sequences [26, 22], hence long-range base pairs in particular must be confirmed independently. We envision a workflow to determine the structure ensembles of an arbitrarily long RNA molecule that begins with DMS-MaPseq [29]. The DMS reactivities would be used [33] to predict two initial models of the structure: one with a limit to the base pair length (for short-range pairs), the other without (for long-range pairs). Sections of the RNA with potential long-range pairs would be flagged from the long-range model and from regions of the short-range model that disagreed with the DMS reactivities (as in Figure 6d). Then, SEARCH-MaP/SEISMIC-RNA could be used to validate, quantify, and refine the potential long-range base pairs; and other methods such as M2-seq [35] to do likewise for short-range base pairs. This integrated workflow could characterize the secondary structures of RNA molecules that have evaded existing methods (e.g. messenger RNAs [21]) as well provide much-needed benchmarks for secondary structure prediction algorithms [25].

In this study, we focused on the genomes of coronaviruses, specifically long-range base pairs involving the frameshift stimulating element (FSE). Long-range base pairs implicated in frameshifting also occur in several plant viruses of the family *Tombusviridae* [65, 66, 67]. However, in *Tombusviridae* species, the frameshift pseudoknots themselves are made of long-range base pairs; in coronaviruses, the pseudoknots are local structures [46, 47, 48, 39] and (at least in SARS-CoV-2) compete with long-range base pairs. Consequently, the long-range base pairs are necessary for frameshifting in *Tombusviridae* species [65, 66, 67] but dispensable in coronaviruses: even the 80-90 nt core FSE of SARS-CoV-2 has stimulated 15-40% of ribosomes to frameshift in dual luciferase constructs [38, 68, 39, 69, 70, 51]. Surprisingly, frameshifting has appeared to be nearly twice as frequent (50-70%) in live SARS-CoV-2 [71, 72, 73]; whether this discrepancy is due to long-range base-pairing, methodological artifacts, or *trans* factors [74] is unknown [75].

If, how, and why the long-range base pairs affect frameshifting in coronaviruses are open questions. For *Tombusviridae*, one study [65] suggested that the long-range stem regulates viral RNA synthesis by negative feedback: without RNA polymerase, the long-range stem would form and stimulate frameshifting to produce polymerase, which would then unwind the long-range stem while replicating the genome. However, this mechanism seems implausible in coronaviruses, where RNA synthesis and translation occur in separate subcellular compartments (the double-membrane vesicles and the cytosol, respectively) [76]. Another study on *Tombusviridae* [67] hypothesized that after the ribosome has frameshifted, long-range stems destabilize the FSE so the ribosome can unwind it and continue translating. As the long-range base pairs in SARS-CoV-2 do compete with the pseudoknot, they might also have this role, which – for coronaviruses – could not be strictly necessary for frameshifting. One study [72] of translation in SARS-CoV-2 at different time points measured frameshifting around 20% at 4 hours post infection but 60-80% at 12-36 hours. This result is consistent with a previous hypothesis [77] that coronaviruses use frameshifting to time protein synthesis: first translating ORF1a to suppress the immune system, then translating ORF1b containing the RNA polymerase. We surmise the long-range base pairs would form in virions and persist when the virus released its genome into a host cell, where they would initially suppress frameshifting. Once host protein synthesis had been inhibited and the double-membrane vesicles formed, a signal specific to the cytosol would disassemble the long-range base pairs so that frameshifting could occur efficiently and produce the replication machinery from ORF1b. The long-range base pairs would form in viral progeny but not in genomic RNA released into the cytosol for translation, so that more ORF1b could be translated. This possible role of long-range base pairs in the coronaviral life cycle could be tested by probing the RNA structure in subcellular compartments and virions, identifying cytosolic factors that could disassemble the long-range base pairs, and quantifying how they affect frameshifting in the context of a live coronavirus.

Future studies could also expand the scope of SEARCH-MaP and SEISMIC-RNA. While all SEARCH-MaP experiments in this study were performed *in vitro*, the method would likely also be feasible *in cellulo*: DMS-MaPseq can detect ASOs binding to RNAs within cells [78]. The main challenges would likely involve optimizing the ASO probes and transfection protocols to maximize the signal while minimizing unwanted side effects such as immunogenicity. SEARCH-MaP can screen an entire transcript (as in Figure 3), but scaling up to an entire transcriptome could prove challenging. One strategy for probing many RNAs simultaneously could involve adding a pool of ASOs – with no more than one ASO capable of binding each RNA – rather than one ASO at a time. In this manner, a similar number of samples would be needed to search all RNAs as would be needed for the longest RNA. Distinguishing direct from indirect base pairing is another area for development: if segment Q could base-pair with either P or R, then blocking P could perturb R (and vice versa) as a consequence of perturbing Q, even though P and R could not base-pair directly. A solution could be to first block Q with one ASO; then, if blocking P with another ASO caused no change in R (and vice versa), it would suggest that they could only interact indirectly (through Q).

We imagine that SEARCH-MaP and SEISMIC-RNA will make it practical to determine accurate secondary structure ensembles of entire messenger, long non-coding, and viral RNAs. Collected in a database of long RNA structures, these results would facilitate subsequent efforts to predict RNA structures and benchmark algorithms, culminating in a real “AlphaFold for RNA” [14] in the hands of every biologist.

## Methods

### Validation of SEISMIC-RNA

SEISMIC-RNA v0.18 was used to simulate and process mutational profiling data for the validation tests, using the script https://github.com/rouskinlab/search-map/tree/main/Compute/validation/run.py. Briefly, 12 reference sequences of 280 nt, secondary structures, and ground-truth mutation rates were generated with SEISMIC-RNA. For each set of ground-truth cluster proportions, nine datasets were simulated with target numbers of 2, 5, 10, 20, 50, 100, 200, 500, and 1,000 thousand reads. Note that the actual number of reads used for clustering (after masking with default parameters) was 50-100% of the target number (e.g. a 20,000-read target could have 10,000-20,000 reads). Each dataset was clustered up to a maximum of 5 clusters; the optimal number of clusters was that which minimized the Bayesian information criterion [30]. After clustering, the number of simulations yielding each number of clusters and the accuracies of the cluster proportions and mutation rates were graphed using the Jupyter notebook https://github.com/rouskinlab/search-map/tree/main/Compute/validation/analyze.ipynb.

### SEARCH-MaP of *E. coli* ribosomes

#### Growth and lysis of *E. coli*

Stellar Chemically Competent Cells (Takara) transformed with an ampicillin resistance plasmid were inoculated into 7 ml of LB Broth (Lennox) containing 100 *µ*g/ml of ampicillin (Sigma-Aldrich). The culture was incubated at 37°C while shaking at 180 rpm for 18.5 hr. OD_600_ of 1 ml was measured using a NanoDrop One cuvette reader. The remaining 6 ml was spun at 21,000 x g. The pellet was resuspended in 1,158 *µ*l of 50 mM Tris-HCl pH 7.4 (Invitrogen) and lysed by adding 1.16 *µ*l of NEBExpress T4 Lysozyme (New England Biolabs) and incubating at room temperature on a tube rotator at 20 rpm for 5 min. The lysate was centrifuged at 16,000 x g at 4°C for 10 min, and the supernatant collected and kept on ice.

#### DMS treatment of *E. coli* lysate

RNA concentration in the supernatant was estimated by purifying 50 *µ*l with an RNA Clean & Concentrator-5 kit (Zymo Research) according to the manufacturer’s protocol, eluting in 20 *µ*l of nuclease-free water (Fisher Bioreagents), and measuring with a NanoDrop One (Thermo Fisher Scientific). Supernatant containing an estimated 1.5 pmol (2.2 *µ*g) of rRNA was mixed with 300 mM Tris-HCl pH 7.4 (Invitrogen), 6 mM magnesium chloride (Invitrogen), and optionally 750 nM ASO (Integrated DNA Technologies) in 98.1 *µ*l; and incubated at 37°C for 20 min to allow the ASO to bind. 1.9 *µ*l DMS (Sigma-Aldrich) was added (to 200 mM) and shaken at 500 rpm in a ThermoMixer C (Eppendorf) at 37°C for 5 min. To quench, 20 *µ*l of beta-mercaptoethanol (Sigma-Aldrich) was added and mixed thoroughly. DMS-modified RNA was purified using an RNA Clean & Concentrator-5 kit (Zymo Research) according to the manufacturer’s protocol, eluted in 10 *µ*l of nuclease-free water (Fisher Bioreagents), and measured with a NanoDrop One (Thermo Fisher Scientific). ASOs and genomic DNA were removed using 0.5 *µ*l TURBO DNase and 1X TURBO DNase Buffer (Invitrogen) in 25 *µ*l at 37°C for 30 min. RNA was purified again using an RNA Clean & Concentrator-5 kit (Zymo Research) and eluted in 10 *µ*l of nuclease-free water (Fisher Bioreagents).

#### Library generation for *E. coli* RNA

100 ng of DMS-modified RNA was prepared for sequencing using the NEBNext UltraExpress RNA Library Prep Kit (New England Biolabs) according to the manufacturer’s protocol for use with NEBNext rRNA Depletion Kits, with the following modifications. The rRNA depletion steps were skipped (probe hybridization, RNase H digestion, and DNase I digestion). Initially, RNA was diluted in nuclease-free water to 25 *µ*l and mixed with 45 *µ*l of RNAClean XP beads (Beckman-Coulter). RNA was fragmented on beads at 94°C for 7 min.

During first strand synthesis, 1 *µ*l of Induro Reverse Transcriptase (New England Biolabs) was used instead of NEBNext UltraExpress First Strand Enzyme Mix; and the reactions were incubated at 25°C for 10 min, 57°C for 30 min, and 95°C for 2 min. AMPure XP Beads (Beckman-Coulter) were used for clean-up after cDNA synthesis. Following adaptor ligation, PCR enrichment was run at half volume (50 *µ*l) for 12 cycles using 20 *µ*l of adaptor-ligated cDNA, 25 *µ*l of NEBNext MSTC High Yield Master Mix, and 5 *µ*l of one pair of indexes from NEBNext Multiplex Oligos for Illumina (Dual Index Primers Set 1). 5 *µ*l of each PCR product was checked on an E-Gel (Invitrogen). The remaining 45 *µ*l were purified using 31.5 *µ*l of AMPure XP Beads, followed by 22.5 *µ*l of 0.1X TE (New England Biolabs) and 18 *µ*l of NEBNext Bead Reconstitution Buffer. Beads were resuspended in 10 *µ*l of 0.1X TE, and 8 *µ*l was transferred to the next step.

DNA concentrations were measured using a Qubit 3.0 Fluorometer (Thermo Fisher Scientific) according to the manufacturer’s protocol. Samples were pooled and sequenced using an iSeq 100 Sequencing System (Illumina) with 2 x 150 bp paired-end reads according to the manufacturer’s protocol.

#### Data analysis of *E. coli* 16S rRNA

Sequencing data were processed with SEISMIC-RNA v0.16 to compute mutation rates, clusters, and correlations between samples using the commands in the shell script https://github.com/rouskinlab/search-map/tree/main/Compute/rrna-lysate-240602/run.sh. Rolling correlations were plotted with the secondary structure from the Comparative RNA Web [15] using the script https://github.com/rouskinlab/search-map/tree/main/Compute/rrna-lysate-240602/plot_genome.py.

### SEARCH-MaP of 2,924 nt SARS-CoV-2 RNA

#### Design of antisense oligonucleotide groups

Antisense oligonucleotides (ASOs, Supplementary Table 1) and their flanking primers (Supplementary Table 2), with constraints on length and melting temperature, were designed using the Python script https://github.com/rouskinlab/search-map/tree/main/Compute/sars2-2924/olaygo.py.

#### Synthesis of 2,924 nt SARS-CoV-2 RNA

A DNA template of the 2,924 nt segment of SARS-CoV-2, including a T7 promoter, was amplified from a previously constructed plasmid [51] in 50 *µ*l using 2X CloneAmp HiFi PCR Premix (Takara Bio) with 250 nM primers TAATACGACTCACTATAGAATAATGAGCTTAGTCCTGTTGCACTACG and TAAATTGCGGACATACTTATCGGCAATTTTGTTACC (Thermo Fisher Scientific); initial denaturation at 98°C for 60 s; 35 cycles of 98°C for 10 s, 65°C for 10 s, and 72°C for 15 s; and final extension at 72°C for 60 s. The 50 *µ*l PCR product with 10 *µ*l of 6X Purple Loading Dye (New England Biolabs) was electrophoresed through a 50 ml gel – 1% SeaKem Agarose (Lonza), 1X tris-acetate-EDTA (Boston BioProducts), and 1X SYBR Safe (Invitrogen) – at 60 V for 60 min. The band at roughly 3 kb was extracted using a Zymoclean Gel DNA Recovery Kit (Zymo Research) according to the manufacturer’s protocol, eluted in 10 *µ*l of nuclease-free water (Fisher Bioreagents), and measured with a NanoDrop (Thermo Fisher Scientific). To increase yield, the gel-extracted DNA was fed into a second round of PCR and gel extraction using the same protocol. Due to remaining contaminants, the DNA was further purified using a DNA Clean & Concentrator-5 kit (Zymo Research) according to the manufacturer’s protocol, eluted in 10 *µ*l of nuclease-free water (Fisher Bioreagents), and measured with a NanoDrop (Thermo Fisher Scientific).

150 ng of DNA template was transcribed using a MEGAscript T7 Transcription Kit (Invitrogen) according to the manufacturer’s protocol, incubating at 37°C for 3 hr. DNA template was then degraded by incubating with 1 *µ*l of TURBO DNase (Invitrogen) at 37°C for 15 min. RNA was purified using an RNA Clean & Concentrator-5 kit (Zymo Research) according to the manufacturer’s protocol, eluted in 20 *µ*l of nuclease-free water (Fisher Bioreagents), and measured with a NanoDrop (Thermo Fisher Scientific).

#### DMS treatment of 2,924 nt SARS-CoV-2 RNA

ASOs were ordered from Integrated DNA Technologies already resuspended to 10 *µ*M in 1X IDTE buffer (10 mM Tris, 0.1 mM EDTA) in a 96-well PCR plate. Each ASO pool was assembled from 25 pmol of each constituent ASO (Supplementary Table 1); volume was adjusted to 12.5 *µ*l by adding TE Buffer – 10 mM Tris (Invitrogen) with 0.1 mM EDTA (Invitrogen). 450 fmol of 2,924 nt SARS-CoV-2 RNA was added to each ASO pool for a total of 13.5 *µ*l in a PCR tube. The tube was heated to 95°C for 60 s to denature the RNA, placed on ice for several minutes, and transferred to a 1.5 ml tube. To refold the RNA, 35 *µ*l of 1.4X refolding buffer comprising 400 mM sodium cacodylate pH 7.2 (Electron Microscopy Sciences) and 6 mM magnesium chloride (Invitrogen) was added, then incubated at 37°C for 25 min. For no-ASO control 1, 12.5 *µ*l of TE Buffer was used instead of an ASO pool. For no-ASO control 2, 12.5 *µ*l of TE Buffer was added after placing on ice and before refolding to confirm the timing of adding TE Buffer would not alter the RNA structure.

RNA was treated with DMS in 50 *µ*l containing 1.5 *µ*l (320 mM) of DMS (Sigma-Aldrich) while shaking at 500 rpm in a ThermoMixer C (Eppendorf) at 37°C for 5 min. To quench, 30 *µ*l of beta-mercaptoethanol (Sigma-Aldrich) was added and mixed thoroughly. RNA was purified using an RNA Clean & Concentrator-5 kit (Zymo Research) according to the manufacturer’s protocol, eluted in 10 *µ*l of nuclease-free water (Fisher Bioreagents), and measured with a NanoDrop (Thermo Fisher Scientific).

ASOs were removed from 4 *µ*l of DMS-modified RNA in 10 *µ*l containing 1 *µ*l of TURBO DNase (Invitrogen) and 1X TURBO DNase Buffer (Invitrogen), incubated at 37°C for 30 min. To stop the reaction, 2 *µ*l of DNase Inactivation Reagent was added and incubated at room temperature for 10 min, mixing several times throughout by flicking. DNase Inactivation Reagent was precipitated by spinning on a benchtop PCR tube centrifuge for 10 min and transferring 4 *µ*l of supernatant to a new tube.

#### Library generation of 2,924 nt SARS-CoV-2 RNA

4 *µ*l RNA was reverse transcribed in 20 *µ*l containing 1X First Strand Buffer (Invitrogen), 500 *µ*M dNTPs (Promega), 5 mM dithiothreitol (Invitrogen), 500 nM FSE primer CTTCGTCCTTTTCTTGGAAGCGACA (Integrated DNA Technologies), 500 nM section-specific reverse primer (Integrated DNA Technologies, Supplementary Table 2), 1 *µ*l of RNaseOUT (Invitrogen), and 1 *µ*l of TGIRT-III enzyme (InGex) at 57°C for 90 min, followed by inactivation at 85°C for 15 min. To degrade the RNA, 1 *µ*l of Hybridase Thermostable RNase H (Lucigen) was added to each tube and incubated at 37°C for 20 min. 1 *µ*l of unpurified RT product was amplified in 12.5 *µ*l using the Advantage HF 2 PCR Kit (Takara Bio) with 1X Advantage 2 PCR Buffer, 1X Advantage-HF 2 dNTP Mix, 1X Advantage-HF 2 Polymerase Mix, 250 nM primers (Integrated DNA Technologies) for either the FSE (CCCTGTGGGTTTTACACTTAAAAAC and CTTCGTCCTTTTCTTGGAAGCGACA) or specific section (Supplementary Table 2); initial denaturation at 94°C for 60 s; 25 cycles of 94°C for 30 s, 60°C for 30 s, and 68°C for 60 s; and final extension at 68°C for 60 s. 5 *µ*l of every amplicon from the same RT product was pooled and then purified using a DNA Clean & Concentrator-5 kit (Zymo Research) according to the manufacturer’s protocol, eluted in 20 *µ*l of nuclease-free water (Fisher Bioreagents), and measured with a NanoDrop (Thermo Fisher Scientific).

200 ng of pooled PCR product was prepared for sequencing using the NEB-Next Ultra II DNA Library Prep Kit for Illumina (New England Biolabs) according to the manufacturer’s protocol with the following modifications. During size selection after adapter ligation, 27.5 *µ*l and 12.5 *µ*l of NEBNext Sample Purification Beads (New England Biolabs) were used in the first and second steps, respectively, to select inserts of 280-300 bp. Indexing PCR was run at half volume (25 *µ*l) for 3 cycles. In lieu of the final bead cleanup, 420 bp inserts were selected using a 2% E-Gel SizeSelect II Agarose Gel (Invitrogen) according to the manufacturer’s protocol. DNA concentrations were measured using a Qubit 3.0 Fluorometer (Thermo Fisher Scientific) according to the manufacturer’s protocol. Samples were pooled and sequenced using an iSeq 100 Sequencing System (Illumina) with 2 x 150 bp paired-end reads according to the manufacturer’s protocol.

#### Data analysis of 2,924 nt SARS-CoV-2 RNA

Sequencing data were processed with SEISMIC-RNA v0.12 and v0.13 to compute mutation rates, clusters, correlations, and secondary structures. Effects of each ASO group (Figure 3b, Supplementary Figures 2 and 3) were computed with the script https://github.com/rouskinlab/search-map/tree/main/Compute/sars2-2924/run-tile.sh. Clustering and structure modeling (Figure 3c and d, Supplementary Figure 4a and b) were performed with the script https://github.com/rouskinlab/search-map/tree/main/Compute/sars2-2924/run-deep.sh. Because some samples contained amplicons that overlapped each other, sequence alignment map (SAM) files were filtered by amplicon using the script https://github.com/rouskinlab/search-map/tree/main/Compute/sars2-2924/filter-deep.py. The fraction of structures containing long-range stems (Supplementary Figure 4c) was determined using the script https://github.com/rouskinlab/search-map/tree/main/Compute/sars2-2924/fraction_folded.py.

### SEARCH-MaP of long-range base pairs in multiple coronaviruses

#### Computational screen for long-range base pairs in coronaviruses

All coronaviruses with reference genomes in the NCBI Reference Sequence Database [54] as of December 2021 were searched for using the following query:

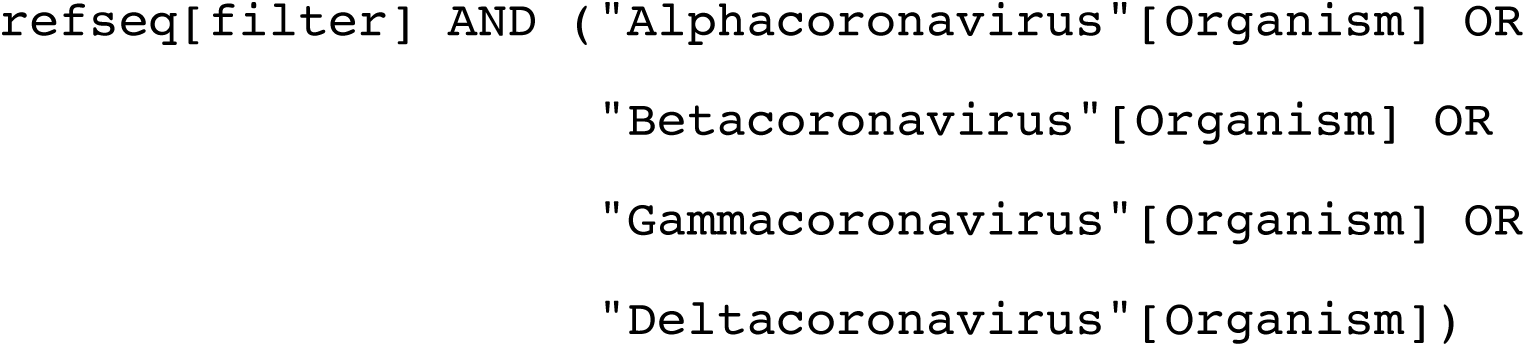

The reference sequences (https://github.com/rouskinlab/search-map/search-map/tree/main/Compute/covs-screen/cov_refseq.fasta) and table of features (https://github.com/rouskinlab/search-map/tree/main/Compute/covs-screen/cov_features.txt) were downloaded and used to locate the slippery site in each genome using a custom Python script (https://github.com/rouskinlab/search-map/tree/main/Compute/covs-screen/extract_long_fse.py). For each genome, up to 100 secondary structure models of the 2,000 nt segment from 100 nt upstream to 1,893 nt downstream of the slippery site (excluding genomes with ambiguous nucleotides in this segment) were generated using Fold v6.3 from RNAstructure [42] via the script https://github.com/rouskinlab/search-map/tree/main/Compute/covs-screen/fold_long_fse.py. The fraction of models in which each base paired with any other base between positions 101 and 250 was calculated using the script https://github.com/rouskinlab/search-map/tree/main/Compute/covs-screen/analyze_interactions.py. Using these fractions, coronaviruses were clustered via the unweighted pair group method with arithmetic mean (UPGMA) and a euclidean distance metric, implemented in Seaborn v0.11 [79] and SciPy v1.7 [80] (Supplementary Figure 6). From each cluster with prominent potential long-range interactions involving the FSE, coronaviruses were manually selected for experimental study.

#### Synthesis of 239 and 1,799 nt coronaviral RNAs

For each selected coronavirus, the 1,799 nt segment from 290 to 1,502 nt downstream of the slippery site was ordered from Twist Bioscience as a gene fragment flanked by the standard 5’ and 3’ adapters CAATCCGCCCTCACTACAACCG and CTACTCTGGCGTCGATGAGGGA, respectively. Gene fragments were resuspended to 10 ng/*µ*l in 10 mM Tris-HCl pH 8 (Invitrogen). Each DNA template for transcription of 1,799 nt RNA segments, including a T7 promoter, was amplified from 0.5 *µ*l (5 ng) of a gene fragment in 20 *µ*l using 2X CloneAmp HiFi PCR Premix (Takara Bio) with 250 nM of each primer TAATACGACTCACTATAGGCAATCCGCCCTCACTACAACCG and TCCCTCATCGACGCCAGAGTAG; initial denaturation at 98°C for 30 s; 30 cycles of 98°C for 10 s, X°C (see Supplementary Table 4) for 10 s, and 72°C for 15 s; and final extension at 72°C for 60 s. DNA templates for transcription of 239 nt RNA segments were amplified using the same procedure but with the forward primers with T7 promoters (F+T7) and reverse primers (R) in Supplementary Table 5. For experiments in which the RNAs were transcribed as a pool of all coronaviruses, all PCR products of the same length (i.e. 239 or 1,799 nt) were pooled, then purified using a DNA Clean & Concentrator-5 kit (Zymo Research) according to the manufacturer’s protocol; concentrations were measured with a NanoDrop (Thermo Fisher Scientific). Otherwise, PCR products were purified individually.

50 ng of DNA template was transcribed using a HiScribe T7 High Yield RNA Synthesis Kit (New England Biolabs) according to the manufacturer’s protocol but at one-quarter volume (5 *µ*l), supplemented with 0.25 *µ*l RNaseOUT (Invitrogen), for 16 hr. DNA template was degraded by incubating with 0.5 *µ*l of TURBO DNase (Invitrogen) at 37°C for 30 min. RNA was purified using an RNA Clean & Concentrator-5 kit (Zymo Research) according to the manufacturer’s protocol, eluted in 50 *µ*l of nuclease-free water (Fisher Bioreagents), and measured with a NanoDrop (Thermo Fisher Scientific).

#### DMS treatment of 239 and 1,799 nt coronaviral RNAs

Antisense oligonucleotides (ASOs) in Supplementary Table 6 were ordered from Integrated DNA Technologies and resuspended to 100 *µ*M in low-EDTA TE buffer: 10 mM Tris pH 7.4 with 0.1 mM EDTA (Integrated DNA Technologies). For each coronavirus, 5 *µ*l of each corresponding ASO (Supplementary Table 6) was pooled; the pool of ASOs was diluted with low-EDTA TE buffer to a final volume of 100 *µ*l, bringing each ASO to 5 *µ*M. 1X refolding buffer comprising 300 mM sodium cacodylate pH 7.2 (Electron Microscopy Sciences) and 6 mM magnesium chloride (Invitrogen) was assembled, then pre-warmed to 37°C.

For already-pooled RNA, 300 ng was diluted in 2.5 *µ*l of nuclease-free water (Fisher Bioreagents) in a PCR tube, heated to 95°C for 1 min to denature, chilled on ice for 3 min, added to 95 *µ*l of pre-warmed refolding buffer, and incubated at 37°C for 20 min to refold. For individually transcribed RNA, 1 pmol was mixed with 10 *µ*l of either low-EDTA TE buffer (for probing without ASOs) or the ASO pool for the corresponding coronavirus (for probing with ASOs) in a PCR tube, heated to 95°C for 1 min to denature the RNA, chilled on ice for 3 min, added to pre-warmed refolding buffer for a total volume of 100 *µ*l, and incubated at 37°C for 20 min to refold the RNA (possibly with ASOs). Subsequently, equimolar amounts of all refolded RNAs were combined into one 97 *µ*l pool in a 1.5 ml tube.

RNA was treated with DMS (Sigma-Aldrich) – 2.5 *µ*l (260 mM) for RNAs transcribed as pools or 3 *µ*l (320 mM) for RNAs pooled after transcription – in 100 *µ*l while shaking at 800 rpm in a ThermoMixer C (Eppendorf) at 37°C for 5 min. To quench, 60 *µ*l of beta-mercaptoethanol (Sigma-Aldrich) was added and mixed thoroughly. DMS-modified RNA was purified using an RNA Clean & Concentrator-5 kit (Zymo Research) according to the manufacturer’s protocol, eluted in 16 *µ*l of nuclease-free water (Fisher Bioreagents), and measured with a NanoDrop (Thermo Fisher Scientific). If added, ASOs were then degraded in

50 *µ*l containing 1X TURBO DNase Buffer (Invitrogen) and 1 *µ*l of TURBO DNase Enzyme (Invitrogen) at 37°C for 30 min; RNA was purified with an RNA Clean & Concentrator-5 kit (Zymo Research) according to the manufacturer’s protocol, eluted in 16 *µ*l of nuclease-free water (Fisher Bioreagents), and measured with a NanoDrop (Thermo Fisher Scientific).

#### Sequencing library generation of 239 and 1,799 nt coronaviral RNAs

100 ng of DMS-modified RNA was prepared for sequencing using the xGen Broad-Range RNA Library Preparation Kit (Integrated DNA Technologies) according to the manufacturer’s protocol, with the following modifications. During fragmentation, 8 *µ*l of RNA was combined with 1 *µ*l of Reagent F1, 4 *µ*l of Reagent F3, and 2 *µ*l of Reagent F2. For reverse transcription, 1 *µ*l of Enzyme R1, 2 *µ*l of TGIRT-III enzyme (InGex), and 1 *µ*l of 100 mM dithiothreitol (Invitrogen) was used instead of the reaction mix, then incubated at room temperature for 30 minutes before adding 2 *µ*l of Reagent F2. Reverse transcription was stopped by adding 1 *µ*l of 4 M sodium hydroxide (Fluka), heating to 95°C for 3 min, chilling at 4°C, then neutralizing with 1*µ*l of 4 M hydrochloric acid. Instead of a bead cleanup after the final PCR, unpurified PCR products with 6X DNA loading dye (Invitrogen) were elecrophoresed through an 8% polyacrylamide Tris-borate-EDTA (TBE) gel (Invitrogen) at 180 V for 55 min. The gel was stained with SYBR Gold (Invitrogen); the section between 250 and 500 bp was excised and placed in a 0.5 ml tube with a hole punctured in the bottom by an 18-gauge needle (BD Biosciences), which was nested inside a 1.5 ml tube and centrifuged at 21,300 x g for 1 min to crush the gel slice into the larger tube. Crushed gel pieces were suspended in 500 *µ*l of 300 mM sodium chloride (Boston Bioproducts), shaken at 1,500 rpm in a ThermoMixer C (Eppendorf) at 70°C for 20 min, and centrifuged at 21,300 x g through a 0.22 *µ*m Costar Spin-X filter column to remove the gel pieces. Filtrate was mixed with 600 *µ*l isopropanol (Sigma-Aldrich) and 3 *µ*l GlycoBlue Coprecipitant (Invitrogen), vortexed briefly, and stored at −20°C overnight. DNA was then pelleted by centrifugation at 4°C at 18,200 x g for 45 min. The supernatant was aspirated, and the pellet was washed with 1 ml of ice-cold 70% ethanol (Sigma-Aldritch), resuspended in 15 *µ*l nuclease-free water (Fisher Bioreagents), and quantified using the 1X dsDNA High Sensitivity Assay Kit for the Qubit 3.0 Fluorometer (Thermo Fisher Scientific) according to the manufacturer’s protocol. Samples were pooled and sequenced using an iSeq 100 Sequencing System (Illumina) with 2 x 150 bp paired-end reads according to the manufacturer’s protocol.

#### Data analysis of 239 and 1,799 nt coronaviral RNAs

Sequencing data were processed with SEISMIC-RNA v0.11 and v0.12 to compute mutation rates, correlations between samples, and secondary structure models using the commands in the shell script https://github.com/rouskinlab/search-map/tree/main/Compute/covs-1799/run.sh. For the 239 and 1,799 nt RNAs that had been pooled during transcription, the two replicates for each coronavirus for each length were confirmed to give similar results, then merged before comparing the 239 and 1,799 nt RNAs to each other. For the comparison of RNAs with and without ASOs, the no-ASO samples that had been transcribed individually were confirmed to give similar results to those transcribed as a pool; then, all no-ASO samples were pooled before comparing to samples with ASOs. For each coronavirus, the DMS reactivities of the combined no-ASO samples were used to model up to 20 secondary structures of the 1,799 nt segment using Fold from RNAstructure v6.3 [42]. Structure models were checked manually for correspondence with the rolling correlation between the +ASO and no-ASO conditions; the minimum free energy structure was chosen for every coronavirus except for transmissible gastroenteritis virus, in which the first sub-optimal structure – but not the minimum free energy structure – contained long-range base pairs supported by the rolling correlation. Rolling correlations between +ASO and no-ASO conditions superimposed on secondary structure models (Figure 5) were graphed using the Python script https://github.com/rouskinlab/search-map/tree/main/Compute/util/pairs_vs_correl.py.

### SEARCH-MaP of 1,799 nt SARS-CoV-2 RNA

#### RNA synthesis of 1,799 nt SARS-CoV-2 RNA

A DNA template for transcription, including a T7 promoter, was amplified from the 1,799 bp gene fragment of SARS-CoV-2 (Twist Bioscience) as described above but with primers TAATACGACTCACTATAGGTACTGGTCAGGCAATAACAGTTACAC and GACCCCATTTATTAAATGGAAAACCAGCTG (Integrated DNA Technologies), an annealing temperature of 65°C, and an extension time of 10 s; eluted in 18 *µ*l of 10 mM Tris-HCl pH 8 (Invitrogen); and measured with a NanoDrop One (Thermo Fisher Scientific). 100 ng of DNA template was transcribed using a HiScribe T7 High Yield RNA Synthesis Kit (New England Biolabs) according to the manufacturer’s protocol for 11 hr. DNA template was degraded by incubating with 1 *µ*l of TURBO DNase (Invitrogen) at 37°C for 30 min. RNA was purified using an RNA Clean & Concentrator-25 kit (Zymo Research) according to the manufacturer’s protocol, eluted in 50 *µ*l of nuclease-free water (Fisher Bioreagents), and measured with a NanoDrop One (Thermo Fisher Scientific).

#### DMS treatment of 1,799 nt SARS-CoV-2 RNA

1.15X refolding buffer comprising 345 mM sodium cacodylate pH 7.2 (Electron Microscopy Sciences) and 7 mM magnesium chloride (Invitrogen) was assembled and pre-warmed to 37°C. 1 pmol of RNA was mixed with 100 pmol of each ASO (Integrated DNA Technologies, Supplementary Table 3) in 10 *µ*l total, heated to 95°C for 60 s to denature, chilled on ice for 5-10 min, and added to 87.1 *µ*l of pre-warmed refolding buffer. If no ASO would be added during refolding, then 1 *µ*l of nuclease-free water (Fisher Bioreagents) was added. RNA was incubated at 37°C for 15-20 min to refold. If an ASO would be added during refolding, then 100 pmol (1 *µ*l) of ASO was added. RNA was incubated for another 15 min to allow any newly added ASOs to bind.

RNA was probed in 100 *µ*l containing 1.9 *µ*l (300 mM) DMS (Sigma-Aldrich) while shaking at 500 rpm in a ThermoMixer C (Eppendorf) at 37°C for 5 min. To quench, 20 *µ*l of beta-mercaptoethanol (Sigma-Aldrich) was added and mixed thoroughly. DMS-modified RNA was purified using an RNA Clean & Concentrator-5 kit (Zymo Research) according to the manufacturer’s protocol, eluted in 15 *µ*l of nuclease-free water (Fisher Bioreagents), and measured with a NanoDrop One (Thermo Fisher Scientific).

#### Library generation 1,799 nt SARS-CoV-2 RNA

1 *µ*l of DMS-modified RNA was reverse transcribed in 20 *µ*l using Induro Reverse Transcriptase (New England Biolabs) according to the manufacturer’s protocol with 500 nM of primer CTTCGTCCTTTTCTTGGAAGCGACA (Integrated DNA Technologies) at 57°C for 30 min, followed by inactivation at 95°C for 1 min. 1 *µ*l of unpurified RT product was amplified in 20 *µ*l using Q5 High-Fidelity 2X Master Mix (New England Biolabs) with 500 nM of each primer CCCTGTGGGTTT-TACACTTAAAAAC and CTTCGTCCTTTTCTTGGAAGCGACA (Integrated DNA Technologies); initial denaturation at 98°C for 30 s; 30 cycles of 98°C for 10 s, 65°C for 20 s, and 72°C for 20 s; and final extension at 72°C for 120 s. The PCR product was purified using a DNA Clean & Concentrator-5 kit (Zymo Research) according to the manufacturer’s protocol, eluted in 20 *µ*l of 10 mM Tris-HCl pH 8 (Invitrogen), and measured with a NanoDrop One (Thermo Fisher Scientific).

50-100 ng of purified PCR product was prepared for sequencing using the NEBNext Ultra II DNA Library Prep Kit for Illumina (New England Biolabs) according to the manufacturer’s protocol with the following modifications. All steps were performed at half of the volume specified in the protocol, including reactions, bead cleanups, and washes. During size selection after adapter ligation, 14 *µ*l and 7 *µ*l of SPRIselect Beads (Beckman Coulter) were used in the first and second steps, respectively, to select inserts of 283 bp. Indexing PCR was run with 400 nM of each primer for 4 cycles. After indexing, PCR products were pooled in pairs; in lieu of the final bead cleanup, 405 bp products were selected using a 2% E-Gel SizeSelect II Agarose Gel (Invitrogen) according to the manufacturer’s protocol. DNA concentrations were measured using a Qubit 4 Fluorometer (Thermo Fisher Scientific) according to the manufacturer’s protocol. Samples were pooled and sequenced using a NextSeq 1000 Sequencing System (Illumina) with 2 x 150 bp paired-end reads according to the manufacturer’s protocol.

#### Data analysis of 1,799 nt SARS-CoV-2 RNA

Sequencing data were processed with SEISMIC-RNA v0.11 and v0.12 to compute mutation rates, clusters, and correlations between samples using the commands in the shell script https://github.com/rouskinlab/search-map/tree/main/Compute/sars2-1799/run.sh.

Heatmaps of the reproducibility of clustering between replicates (Supplementary Figure 5) were generated using the Python script https://github.com/rouskinlab/search-map/tree/main/Compute/sars2-1799/compare-clusters.py. After the two replicates were confirmed to give similar clusters, they were pooled for subsequent analyses. Secondary structures with rolling correlations (Figure 4b) were drawn using the Python script https://github.com/rouskinlab/search-map/tree/main/Compute/sars2-1799/draw-structure.py. Alternative structure models (Figure 4c) were selected and created with the help of the Python scripts https://github.com/rouskinlab/search-map/tree/main/Compute/sars2-1799/choose-model-parts.py and https://github.com/rouskinlab/search-map/tree/main/Compute/sars2-1799/make-models.py. Heatmaps of areas under the curve (Figure 4d) were generated using the Python script https://github.com/rouskinlab/search-map/tree/main/Compute/sars2-1799/atlas-plot.py.

### DMS-MaPseq of transmissible gastroenteritis virus in ST cells

#### Cells and Viruses

Transmissible gastroenteritis virus (TGEV, TC-adapted Miller strain, ATCC VR-1740) and ST cells (ATCC CRL-1746) were ordered from American Type Culture Collection (ATCC). ST cells were maintained in Eagle’s Minimum Essential Medium (EMEM, Gibco) supplemented with 10% fetal bovine serum (Gibco), 1% sodium pyruvate (Gibco), and 1% Pen Strep (Gibco) at 37°C with 5% carbon dioxide. For TGEV, the infection medium (IM) comprised EMEM (Gibgo) supplemented with 10% fetal bovine serum (Gibco), 1% sodium pyruvate, and 1 *µ*g/ml of TPCK trypsin (Thermo Fisher Scientific).

#### Production and titering of TGEV

A 150 mm dish was seeded with 1 x 10^7^ ST cells, grown overnight, and washed twice with phosphate-buffered saline (PBS, Gibco). Cells were inoculated with 8 ml of TGEV in IM at a multiplicity of infection (MOI) of 0.1, which was kept on for 60 min with rocking every 15 min. The inoculum was removed, cells were washed twice with PBS, and 26 ml of IM was added. Cells were checked daily for cytopathic effects (CPE); after 5 days, upon development of 80% CPE, the supernatant was centrifuged at 200 x g for 5 min and then filtered through a 0.45 *µ*m filter to harvest TGEV, which was frozen at −80°C in 1 ml aliquots.

Harvested TGEV was titered via tissue culture infectious dose (TCID_50_). Briefly, ST cells were seeded in a poly-L-lysine coated 96-well plate at 4 x 10^4^ cells per well and grown overnight. TGEV was thawed on ice and serially diluted from 10^-1^ to 10^-10^ in IM. Cells were washed once with PBS, and each well was inoculated with 100 *µ*l of either one serial dilution of TGEV (8 replicates per dilution level) or sterile IM (for negative controls). The plate was wrapped in parafilm and incubated at 37°C until CPE appeared. Then, media was aspirated and cells were fixed with 4% paraformaldehyde for 30 min and decanted. 0.5% crystal violet was then added to each well; the plate was rocked for 10 min, submerged in water to remove excess crystal violet, and dried. Wells with CPE were counted and the titer determined using the Spearman-Kärber method.

#### TGEV infection and DMS treatment

Four 150 mm dishes were each seeded with 1 x 10^7^ ST cells, grown overnight, and washed twice with phosphate-buffered saline. Cells were inoculated with 8 ml of TGEV in IM at a multiplicity of infection (MOI) of 2, which was kept on for 60 min with rocking every 15 min. The inoculum was removed, cells were washed twice with PBS, and 26 ml of IM was added.

After 48 hr, media was aspirated. 250 *µ*l of DMS was mixed with 10 ml of IM and immediately added to two plates; the other two received 10 ml IM without DMS. Plates were incubated at 37°C for 5 min. The media was aspirated and replaced with stop solution (30% beta-mercaptoethanol in 1X PBS). Cells were scraped off using a cell scraper and spun down at 3,000 x g for 3 min. The pellet was washed with stop solution, spun down again, washed with 10 ml PBS, dissolved in 3 ml of TRIzol (Invitrogen), and split into 1 ml techincal replicates.

#### RNA purification

200 *µ*l of chloroform was added to each 1 ml techincal replicate, vortexed for 20 s, and rested until the phases separated. Samples were then spun at 18,200 x g for 15 min at 4°C; the aqueous phase transferred to a new tube and mixed with an equal volume of 100% ethanol (Koptec). RNA was purified using a 50 *µ*g Monarch RNA Cleanup Column (New England Biolabs), eluted in 20 *µ*l of nuclease-free water, and quantified with a NanoDrop.

To remove rRNA, 10 *µ*g of total RNA was diluted in 6 *µ*l of nuclease-free water and mixed with 1 *µ*l of anti-rRNA ASOs (Integrated DNA Technologies) and 3 *µ*l of HYBE buffer (200 mM sodium chloride, 100 mM Tris-HCl pH 7.5). The mixture was incubated at 95°C for 2 min and cooled by 0.1°C/s until reaching 45°C. A preheated mixture of 10 *µ*l of RNase H (New England Biolabs) and 2 *µ*l of RNase H Buffer (New England Biolabs) was added and incubated at 45°C for 30 min. RNA was purified using a 10 *µ*g Monarch RNA Cleanup Column (New England Biolabs) and eluted in 42 *µ*l of nuclease-free water.

To remove DNA (including anti-rRNA ASOs), 5 *µ*l of 10X Turbo DNase Buffer (Thermo Fisher) and 3 *µ*l of TURBO RNase (Thermo Fisher) were added and incubated at 37°C for 20 min. RNA was purified using a 10 *µ*g Monarch RNA Cleanup Column (New England Biolabs) and eluted in 10 *µ*l of nuclease-free water.

#### Library generation for the full TGEV genome

RNA was prepared for sequencing using the xGen Broad-Range RNA Library Preparation Kit (Integrated DNA Technologies) according to the manufacturer’s protocol, with the same modifications as described above (5.4.4), notably the substitution of TGIRT-III (InGex) for the kit’s reverse transcriptase. Samples were pooled and sequenced using a NextSeq 1000 Sequencing System (Illumina) with 2 x 150 bp paired-end reads according to the manufacturer’s protocol.

#### Library generation for amplicons

1 *µ*l of rRNA-depleted, DNased RNA was reverse transcribed in 20 *µ*l using Induro Reverse Transcriptase (New England Biolabs) according to the manufacturer’s protocol with 500 nM of primer ACAATTCGTCTTAAGGAATTTACCAATACACGCAA (Integrated DNA Technologies) at 57°C for 30 min, followed by inactivation at 95°C for 1 min. 1 *µ*l of unpurified RT product was amplified in 10 *µ*l using Q5 High-Fidelity 2X Master Mix (New England Biolabs) with 1 *µ*M of each primer, either GCCGCTACAAAGGTAAGTTCGTGCAAATACCAACT and ACAATTCGTCTTAAGGAATTTACCAATACACGCAA or GTGAAAAGTGACATCTATGGTTCTGATTATAAGCAGTA and CTATACCAAGTTGTTTGAAATGGTAACCTGCAGTAACA (Integrated DNA Technologies); initial denaturation at 98°C for 30 s; 30 cycles of 98°C for 5 s, 69°C for 20 s, and 72°C for 15 s; and final extension at 72°C for 120 s. Amplification was confirmed by electrophoresing 1 *µ*l of each PCR product. PCR products for both pairs of primers were pooled and then purified using a DNA Clean & Concentrator-5 kit (Zymo Research) according to the manufacturer’s protocol, eluted in 18 *µ*l of 10 mM Tris-HCl pH 8 (Invitrogen), and measured with a NanoDrop (Thermo Fisher Scientific).

175-225 ng of purified PCR product was prepared for sequencing using the NEBNext Ultra II DNA Library Prep Kit for Illumina (New England Biolabs) according to the manufacturer’s protocol with the following modifications. All steps were performed at half of the volume specified in the protocol, including reactions, bead cleanups, and washes. During size selection after adapter ligation, 14 *µ*l and 7 *µ*l of SPRIselect Beads (Beckman Coulter) were used in the first and second steps, respectively, to select inserts of 295 bp. Indexing PCR was run with 400 nM of each primer for 4 cycles. In lieu of the final bead cleanup, 415 bp products were selected using a 2% E-Gel SizeSelect II Agarose Gel (Invitrogen) according to the manufacturer’s protocol. DNA concentrations were measured using a Qubit 4 Fluorometer (Thermo Fisher Scientific) according to the manufacturer’s protocol. Samples were pooled and sequenced using a NextSeq 1000 Sequencing System (Illumina) with 2 x 150 bp paired-end reads according to the manufacturer’s protocol.

#### Data analysis of transmissible gastroenteritis virus in ST cells

The genomic sequence of this TGEV strain was determined using the script https://github.com/rouskinlab/search-map/tree/main/Compute/tgev-virus/consensus.sh: reads from the untreated sample were aligned to the TGEV reference genome (`NC_038861.1`) using Bowtie 2 [41] and the consensus sequence was determined using Samtools [81]. All reads were processed with SEISMIC-RNA v0.17 to compute mutation rates, correlations between samples, and secondary structure models using the commands in the shell script https://github.com/rouskinlab/search-map/tree/main/Compute/tgev-virus/run.sh. Positions in the untreated sample with mutation rates greater than 0.01 were masked. Replicates were checked for reproducibility and pooled for clustering and structure modeling. A model of short-range base pairs (maximum distance 300 nt) in the TGEV genome was generated from the DMS reactivities using Fold-smp from RNAstructure [42] in five overlapping 10 kb segments, which were merged using the script https://github.com/rouskinlab/search-map/tree/main/Compute/tgev-virus/assemble-tgev-ss.py. Rolling area under the curve superimposed on secondary structure models in Figure 6d was graphed using the script https://github.com/rouskinlab/search-map/tree/main/Compute/tgev-virus/make-figure-6d.py, and in Supplementary Figure 10 using the script https://github.com/rouskinlab/search-map/tree/main/Compute/tgev-virus/plot_genome.py.

## Acknowledgements

We thank Miriam L. Rittenberg and Mateo Valenzuela for assistance with experiments and Daniel Herschlag for advice on the project and the manuscript. This research was supported by National Institute of Allergy and Infectious Diseases grant DP2 AI175475 (S.R.); National Science Foundation Graduate Research Fellowship Program grants 1745302 (M.F.A.), 2140743 (J.A.), and 2141064 (M.F.A.); and National Institute of General Medical Sciences grants T32 GM145407 (J.A.) and R01 GM132899 (D.H.).

## Author Contributions

S.R. and M.F.A. conceived the project. M.F.A. performed the experiments with ribosomes. M.F.A. performed the experiments with SARS-CoV-2 segments. J.A. performed the experiments with other coronavirus segments. J.S.P. performed the experiments with TGEV-infected ST cells. M.F.A. wrote SEISMIC-RNA with contributions from S.L.G., Y.J.M.T., A.A.L., and J.A. M.F.A. benchmarked SEISMIC-RNA. M.F.A. analyzed the data with contributions from J.A. M.F.A. drafted the manuscript. All authors reviewed the manuscript and provided comments.

## Ethics Declarations

The authors declare no competing interests.

## Data Availability

All sequencing data generated in this study have been deposited into the NCBI Short Read Archive under accession code PRJNA1103196.

## Code Availability

Documentation for SEISMIC-RNA, including instructions for installation, is hosted on GitHub Pages: https://rouskinlab.github.io/seismic-rna. Source code for SEISMIC-RNA is available from GitHub: https://github.com/rouskinlab/seismic-rna. Shell scripts for running SEISMIC-RNA, auxiliary scripts for data analysis, supplementary files, and LaTeX source code for this manuscript are also available from GitHub: https://github.com/rouskinlab/search-map.

