## Supplementary Information for "Discovery and Quantification of Long-Range RNA Base Pairs in Coronavirus Genomes with SEARCH-MaP and SEISMIC-RNA"

### Supplementary Figures

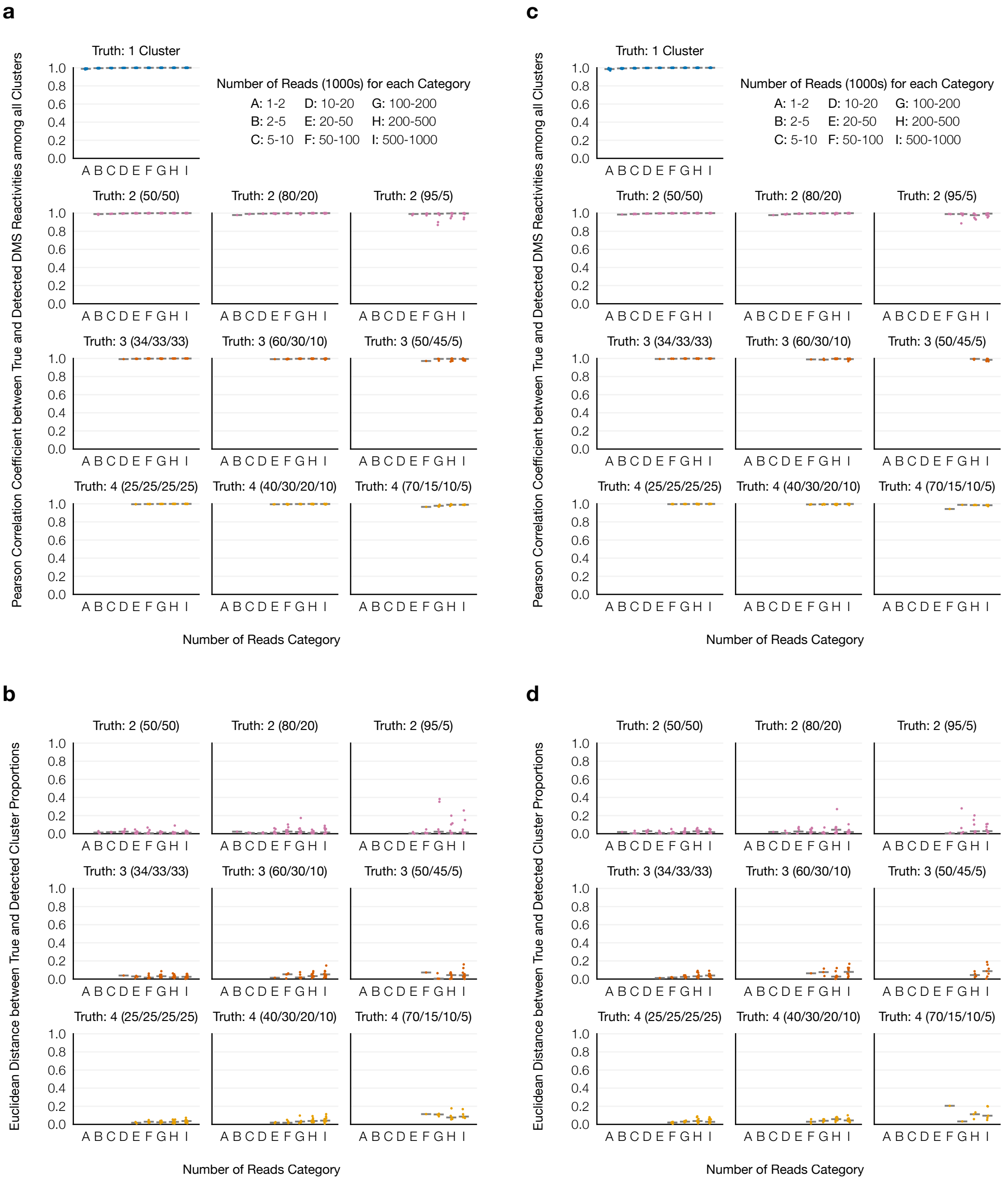


Supplementary Figure 1: **Accuracy of SEISMIC-RNA's clustering algorithm.** Datasets were simulated with one to four "true" alternative structure(s) in three mixing proportions and with 1,000 to 1,000,000 reads (all full-length). For each number of reads, 12 unique 280-nt RNAs were simulated and processed with SEISMIC-RNA. For simulations in which the correct number of clusters was detected, **(a)** the Pearson correlation of the true DMS reactivities versus those calculated by the clustering algorithm and **(b)** the Euclidean distance between the vector of true cluster proportions and those calculated by the clustering algorithm is shown as one point; medians are shown as gray bars. **(c)** and **(d)** Same as (a) and (b), respectively, but reads were simulated with random 5' and 3' ends rather than fully covering the RNA.

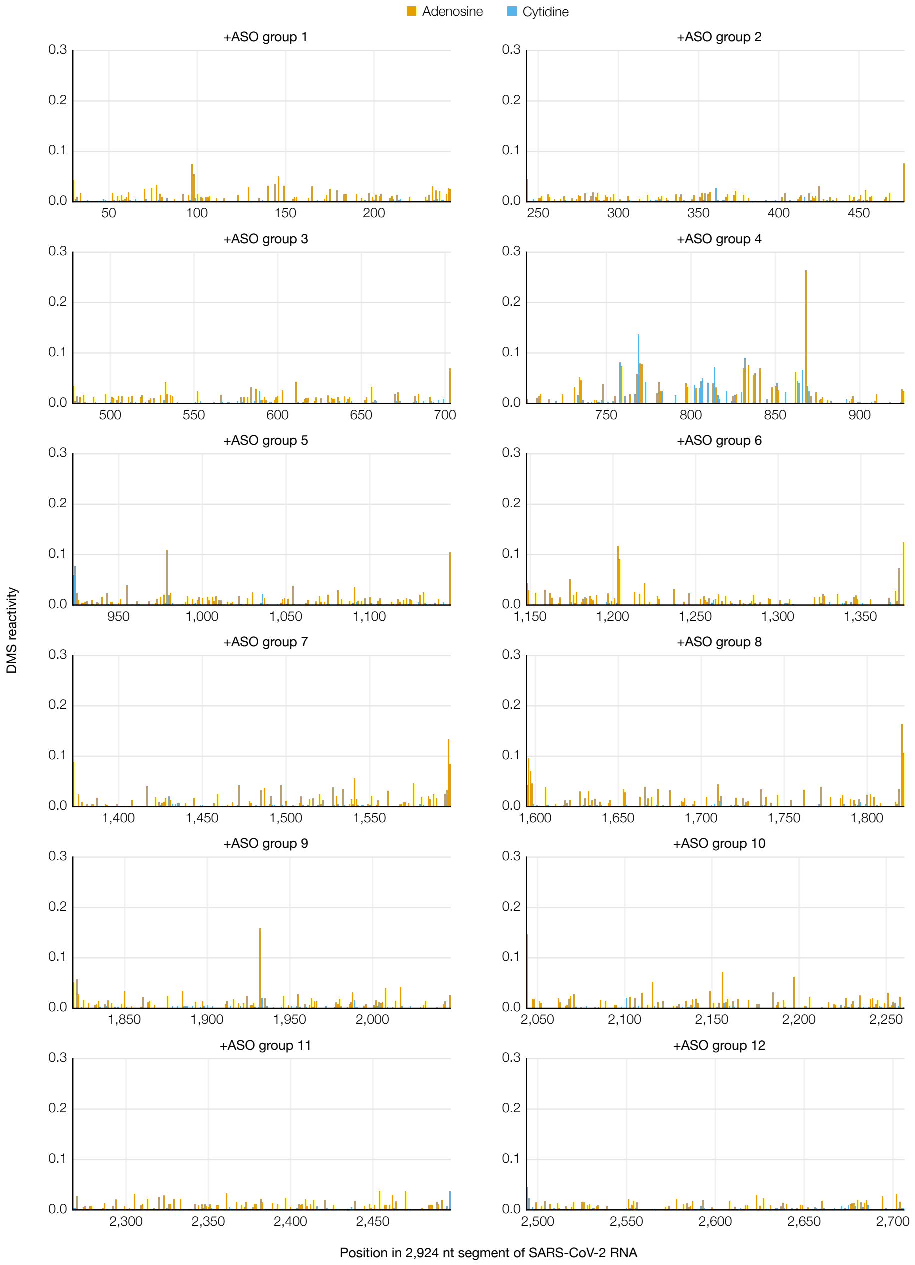


Supplementary Figure 2: **Mutational profile of each ASO target section upon adding the corresponding group of ASOs to the 2,924 nt segment of SARS-CoV-2 genomic RNA.** Positions are colored based on the RNA sequence.

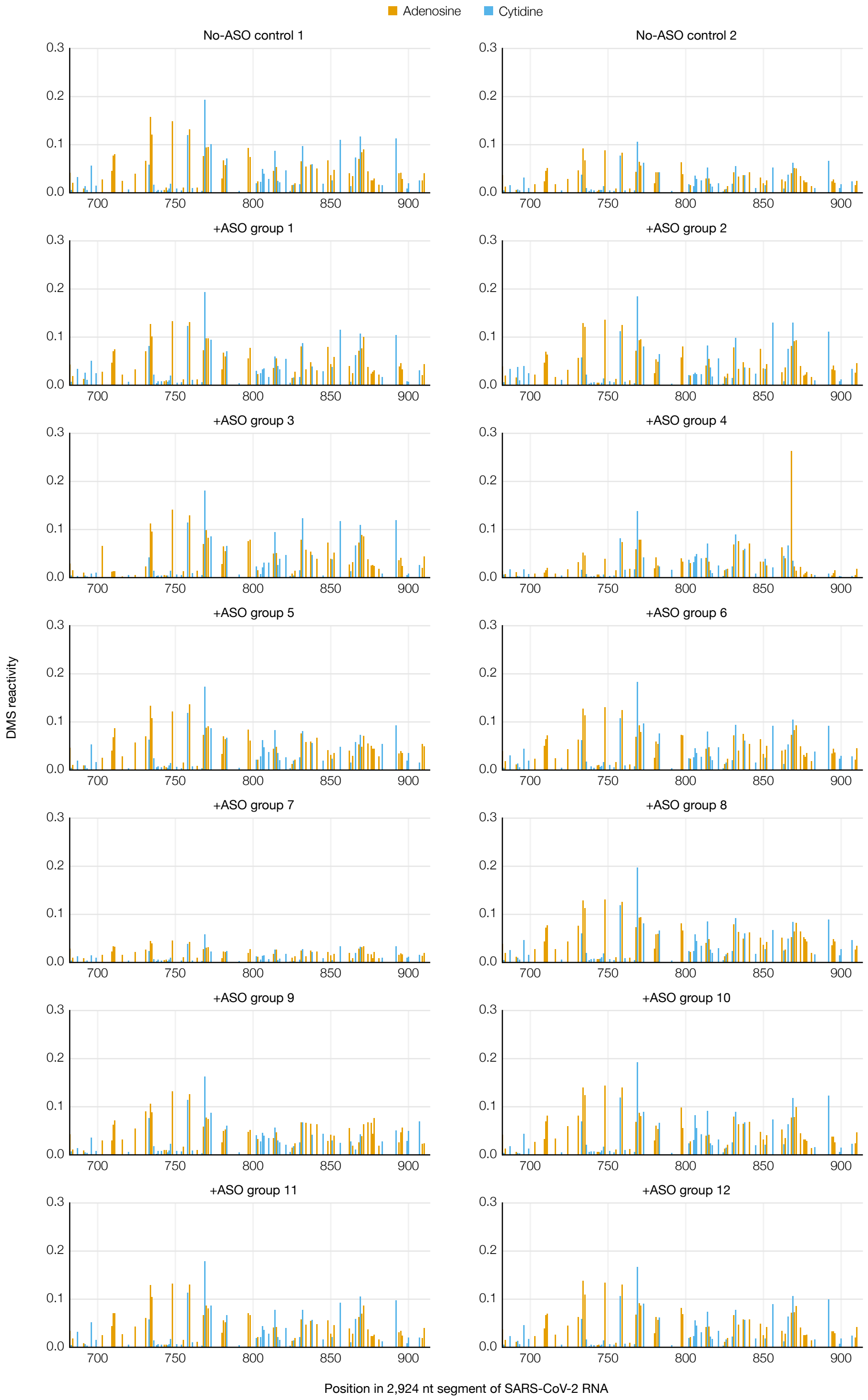


Supplementary Figure 3: **Mutational profiles of the FSE section upon adding each group of ASOs to the 2,924 nt segment of SARS-CoV-2 genomic RNA.** Positions are colored based on the RNA sequence.

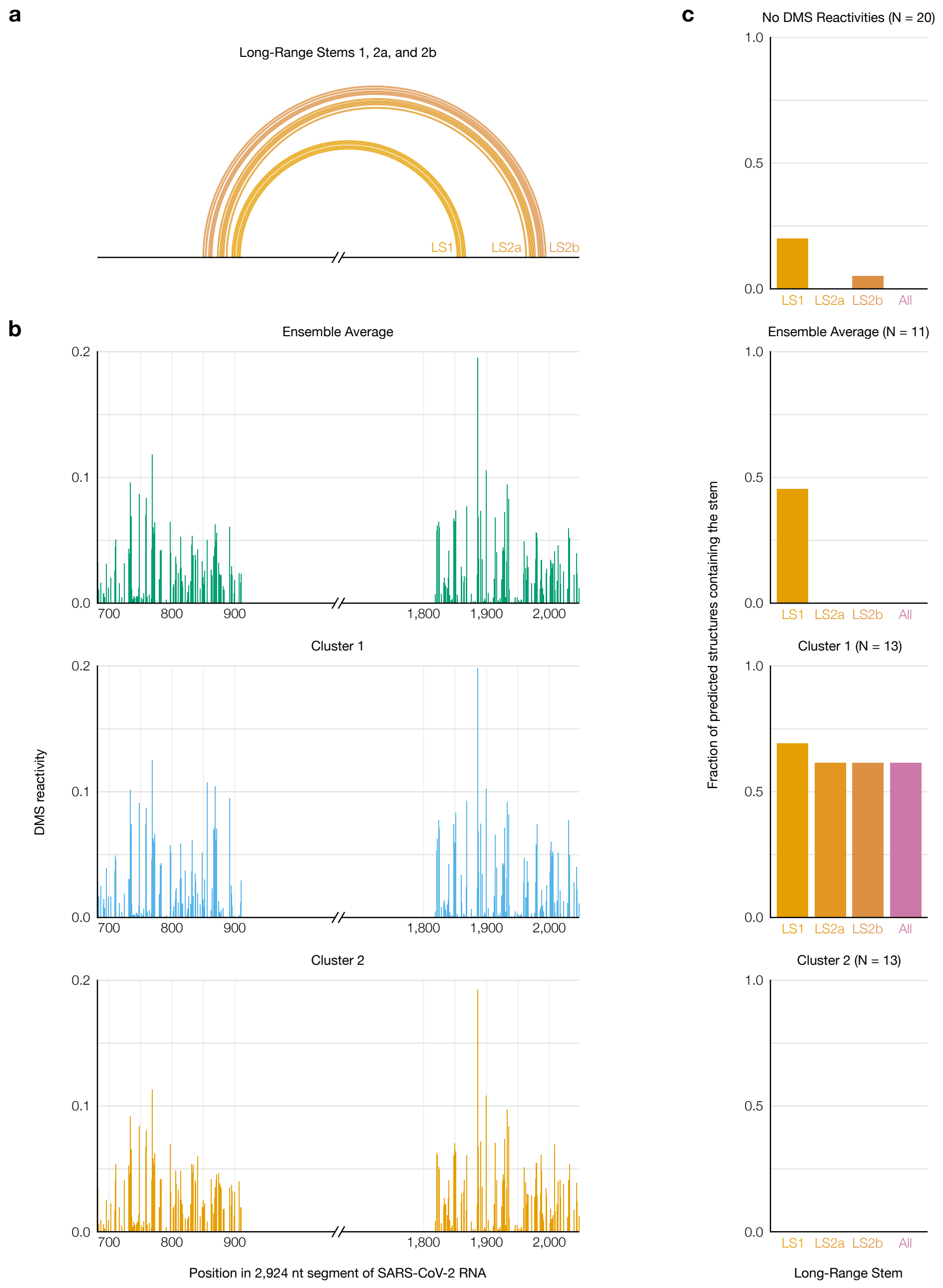


Supplementary Figure 4: **Improved prediction of long-range stems in SARS-CoV-2 using clustered DMS reactivities. (a)** Model of the two inner stems of the FSE-arch [52], denoted long stems (LS) 1 and 2a/b. **(b)** Mutational profiles of the ensemble average and of clusters 1 and 2 on both sides of the FSE-arch. **(c)** For each mutational profile (as well as a purely thermodynamic prediction with no DMS reactivities), the fraction of predicted structures in which each long stem was predicted perfectly (i.e. all base pairs were present). The numbers of predicted structures (N) are indicated.

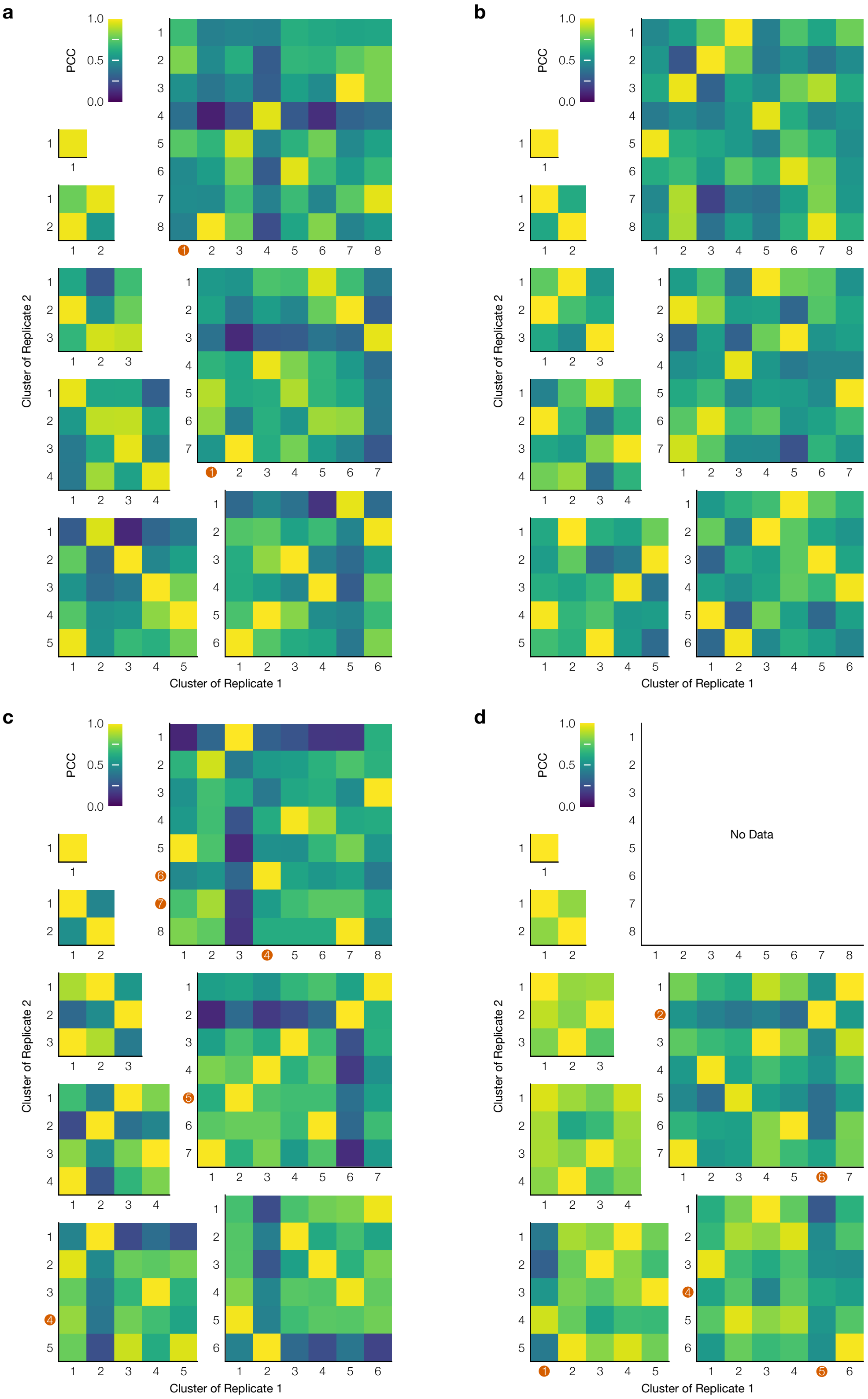


Supplementary Figure 5: **Reproducibility of clustering the SARS-CoV-2 FSE after adding ASOs.** **(a)** Heatmaps of the Pearson correlation coefficient (PCC) between each pair of clusters from two replicates of the 1,799 nt segment of SARS-CoV-2. Each heatmap corresponds to one order (i.e. number of clusters). Clusters are marked with red circles if at least one DMS reactivity exceeded 0.3. **(b)** Same as (a) plus Anti-AS1 ASO. **(c)** Same as (a) plus Anti-PS2-overlap ASO. **(d)** Same as (a) plus Anti-AS1 and Anti-PS2-overlap ASOs.

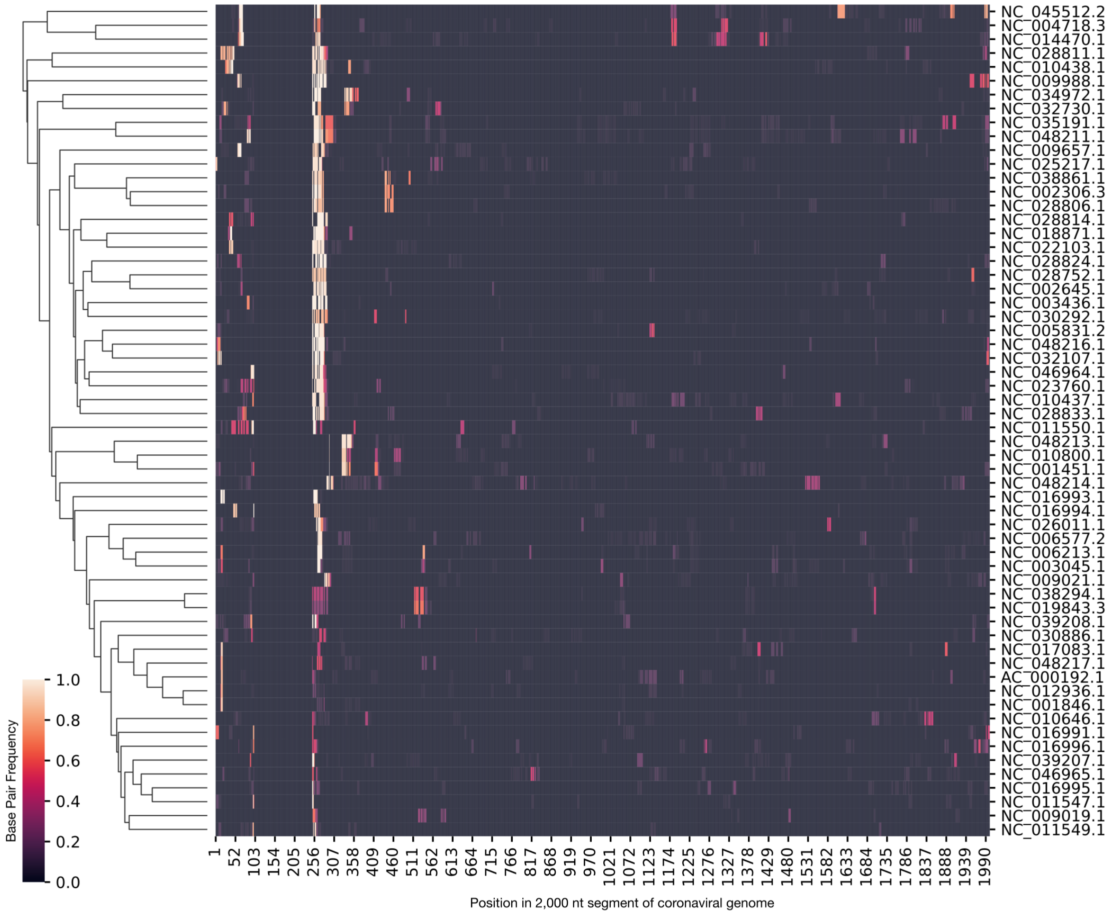


Supplementary Figure 6: **Computational screen of long-range base pairing near the FSE in 60 coronaviruses.** For each 2,000 nt segment of each coronaviral genome, the fraction of predicted structures in which each position outside the range 101-250 base-paired with any position in the range 101-250 is indicated. Genomes are clustered by their base-pairing frequencies. For each genome, the accession number for NCBI [54] is indicated.

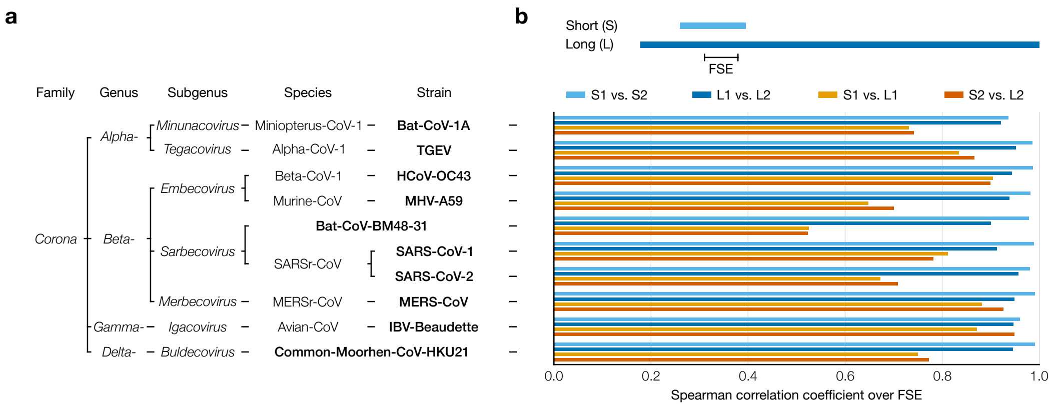


Supplementary Figure 7: **Experimental screen of long-range base pairing near the FSE in 10 coronaviruses.** **(a)** Taxonomy of the ten coronavirus species/strains in this screen; the lowest-level group for each virus is bolded. Bat-CoV-1A: bat coronavirus 1A (NC_010437.1), TGEV: transmissible gastroenteritis virus (NC_038861.1), HCoV-OC43: human coronavirus OC43 (NC_006213.1), MHV-A59: murine hepatitis virus strain A59 (NC_048217.1), Bat-CoV-BM48-31: bat coronavirus BM48-31 (NC_014470.1), SARS-CoV-1: severe acute respiratory syndrome coronavirus 1 (NC_004718.3), SARS-CoV-2: severe acute respiratory syndrome coronavirus 2 (NC_045512.2), MERS-CoV: Middle East respiratory syndrome coronavirus (NC_019843.3), IBV-Beaudette: avian infectious bronchitis virus strain Beaudette (NC_001451.1), Common-Moorhen-CoV-HKU21: common moorhen coronavirus HKU21 (NC_016996.1). **(b)** Spearman correlation coefficients of DMS reactivities over the FSE between replicates 1 and 2 of short (239 nt) and long (1,799 nt) segments of each coronaviral genome.

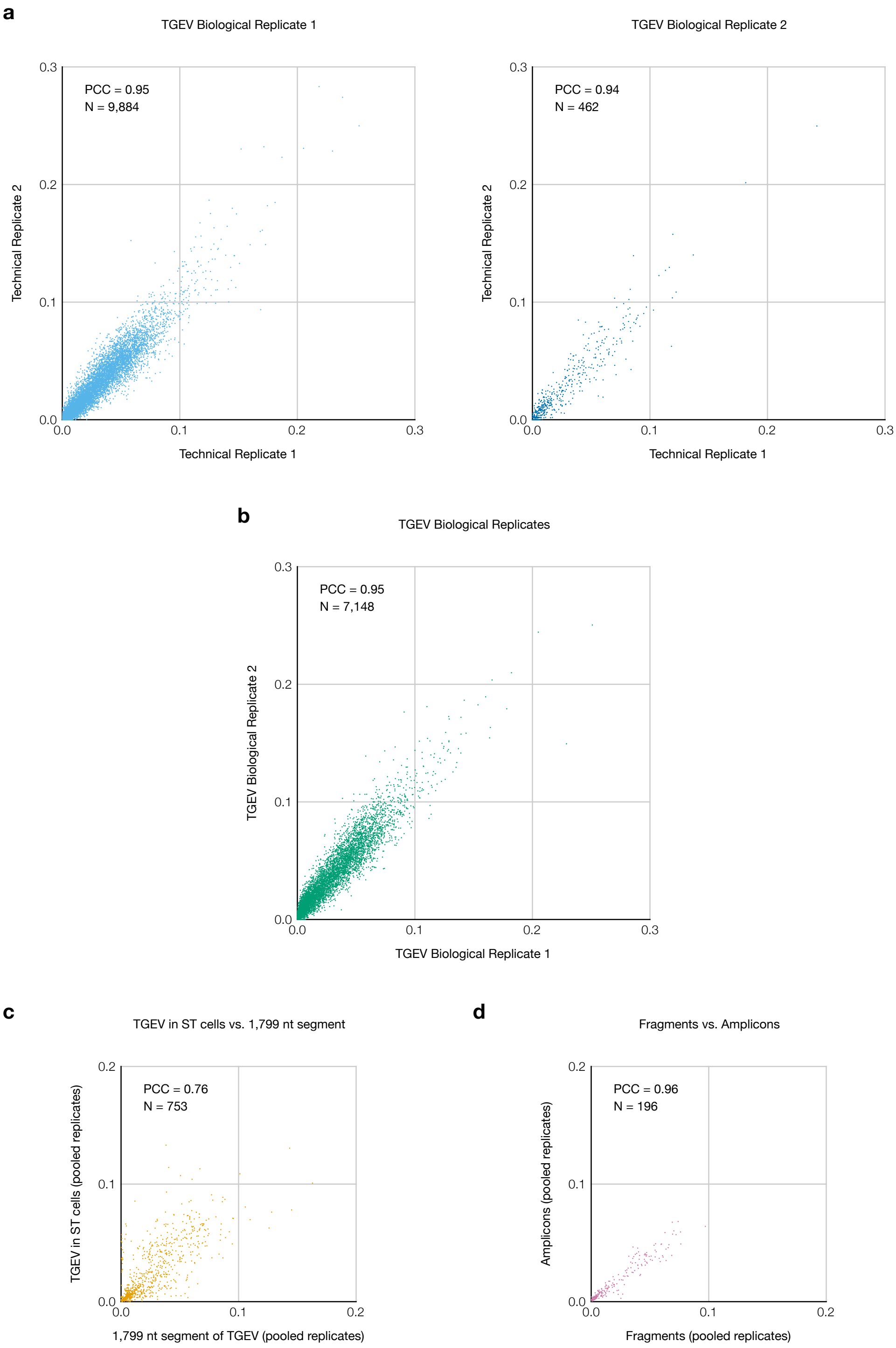


Supplementary Figure 8: **Replicates of TGEV in ST cells and comparison to the 1,799 nt segment. (a)** Comparison of DMS reactivities of the two technical replicates for each biological replicate of TGEV in ST cells. Each point represents one base in the sequence. The number of points (N) and Pearson correlation coefficient (PCC) are indicated for each plot. Two bases with DMS reactivities exceeding 0.3 in both technical replicates of biological replicate 1 are not shown. **(b)** Comparison of DMS reactivities of the two biological replicates (pooled technical replicates). One base with DMS reactivity exceeding 0.3 in biological replicate 1 is not shown. **(c)** DMS reactivities of TGEV in ST cells using random fragmentation versus amplicons (pooled biological replicates). **(d)** DMS reactivities of TGEV in ST cells (pooled biological replicates) versus the 1,799 nt segment.

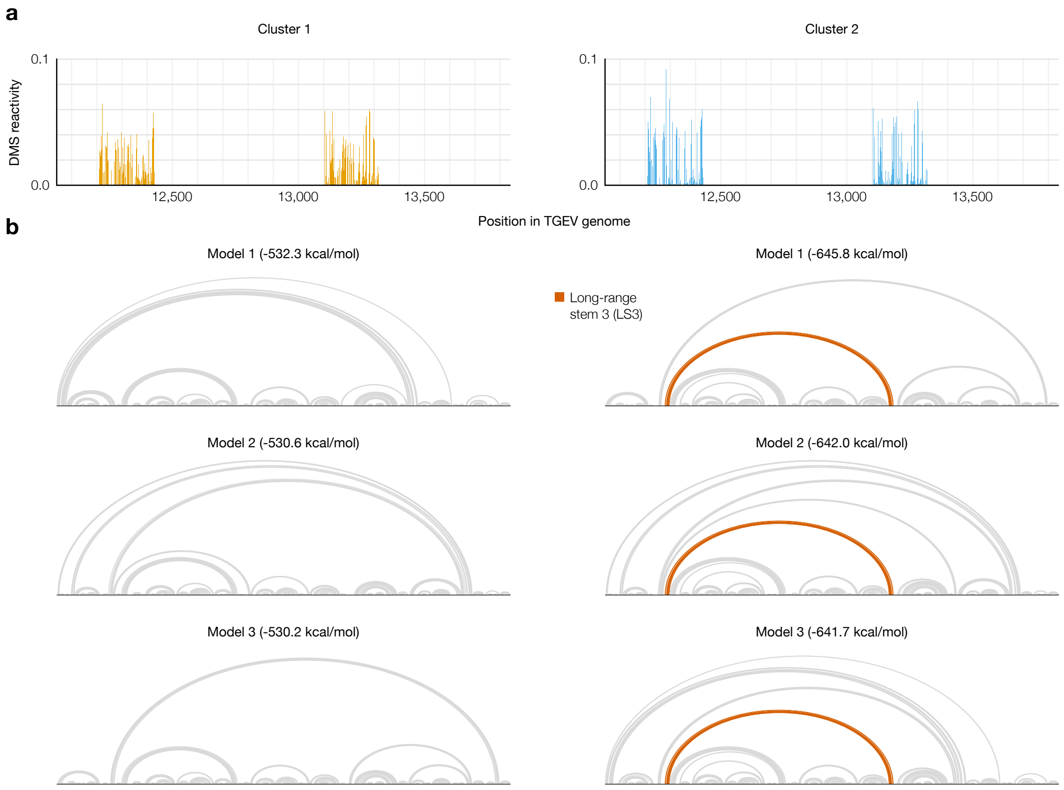


Supplementary Figure 9: **Alternative structures on both sides of the long-range base pairs in TGEV. (a)** DMS reactivities of clusters 1 and 2 on both sides of the long-range base pairs in TGEV, from amplicon samples. **(b)** Three lowest-energy structure models of the 1,799 nt segment (positions 12,042-13,840) based on the DMS reactivities of each cluster. Long-range stem 3 (LS3) is highlighted when it appears in a model. Structures were drawn with VARNA [82].

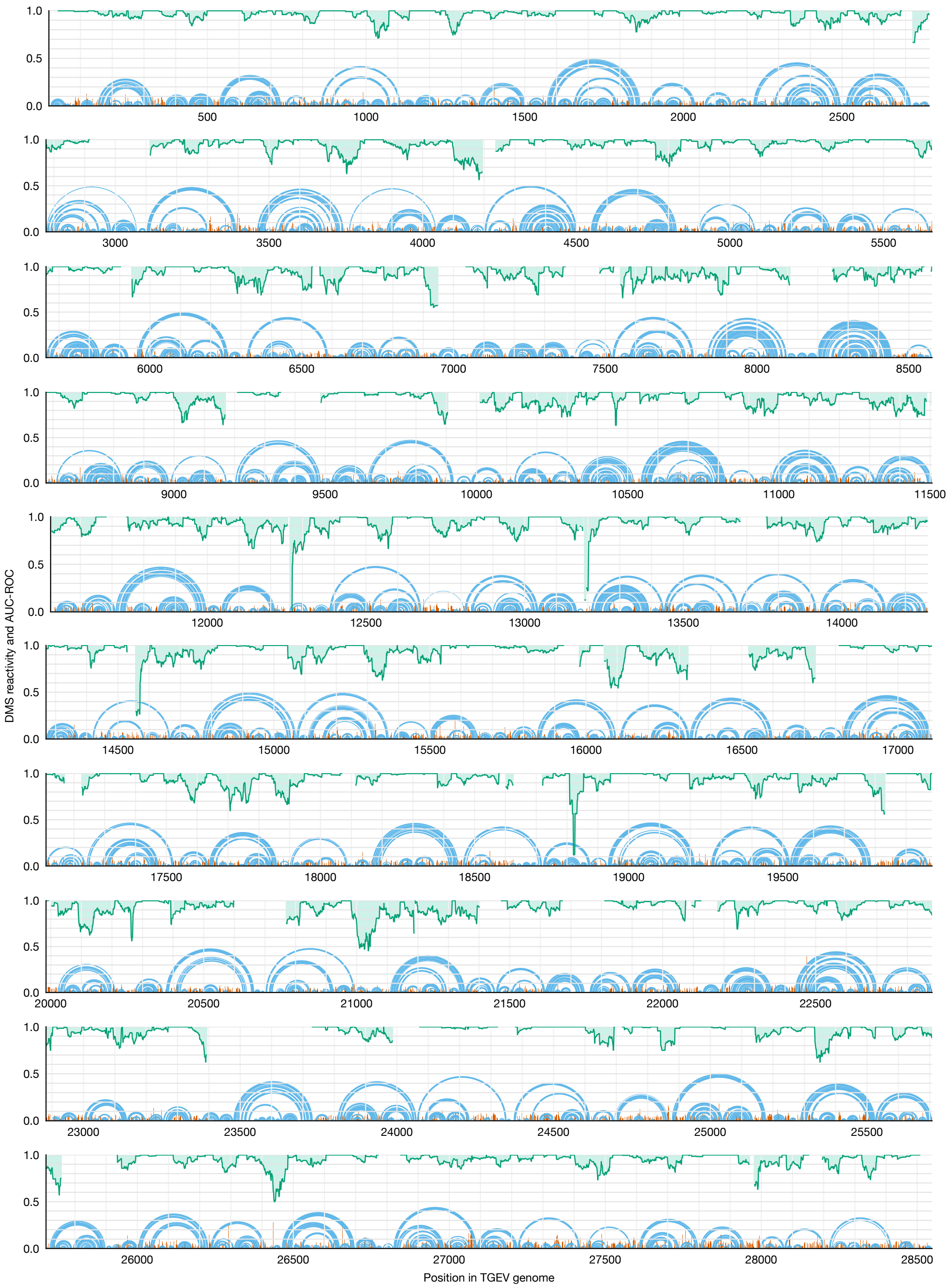


Supplementary Figure 10: **Short-range base pairs across the full TGEV genome.** Model of the secondary structure of the entire TGEV genome with a maximum distance of 300 nt between paired bases (blue). DMS reactivities used to generate the model are shown in red. Rolling (window = 45 nt) area under the receiver operating characteristic curve (AUC-ROC), measuring how well the secondary structure model fits the DMS reactivities, is shown in green.

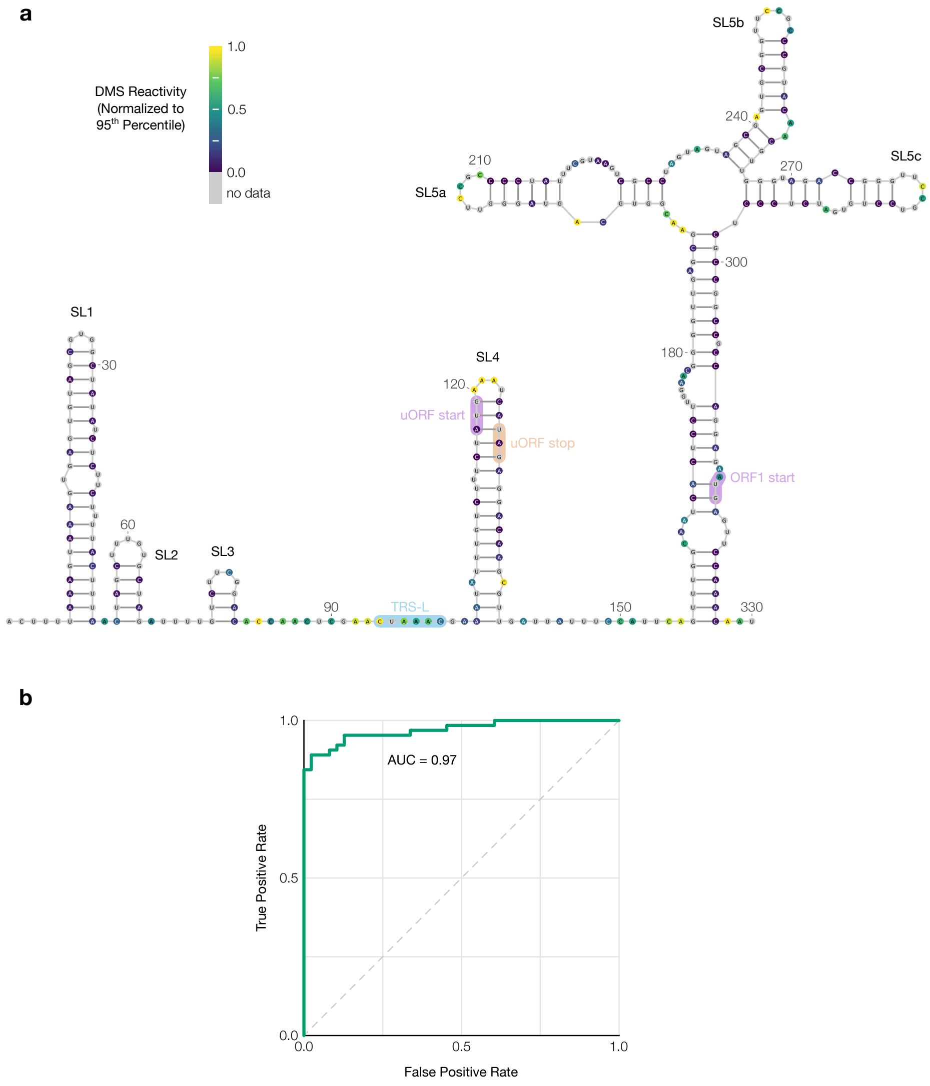


Supplementary Figure 11: **Secondary structure of the TGEV 5' UTR. (a)** Model of the secondary structure of the first 330 nt of the TGEV genome, based on DMS reactivities in infected ST cells normalized to the 95th percentile. Bases are colored by DMS reactivity. The model includes the conserved stem loops SL1, SL2, SL3, SL4, SL5a, SL5b, and SL5c [10]. The leader transcription regulatory sequence (TRS-L) [83], upstream open reading frame (uORF) [84], and start codon of ORF1 are also labeled. The model was drawn using VARNA [82]. **(b)** Receiver operating characteristic curve showing agreement between the DMS reactivities and the secondary structure model; the area under the curve (AUC) is indicated.

### Supplementary Methods

#### Correcting observer bias due to drop-out of reads

Let $N$ reads from $K$ clusters align to a reference sequence of length $L$. Let the proportion of reads whose 5' and 3' ends align, respectively, to coordinates $a$ and $b$ ($1 \leq a \leq b \leq L$) be $\eta_{ab}$ (assuming these proportions are equal for all clusters). Let the mutation rate of base $j$ ($1 \leq j \leq L$) in cluster $k$ ($1 \leq k \leq K$) be $\mu_{jk}$. Let the proportion of cluster $k$ in the ensemble be $\pi_k$. To express these quantities as probabilities, let $C_k$ be the event that a read comes from cluster $k$; let $E_{ab}$ be the event that a read aligns with 5' and 3' coordinates $a$ and $b$, respectively; let $S_j$ be the event that a read contains position $j$ (i.e. its alignment coordinates $a$ and $b$ satisfy $1 \leq a \leq j \leq b \leq L$); let $M_j$ be the event that a read has a mutation at position $j$; and let $G_g$ be the event that a read has no two mutations separated by fewer than $g$ non-mutated bases.

##### Deriving mutation rates of reads with no two mutations too close

In terms of these events, the total mutation rates ($\mu_{jk}$) are $P(M_j|S_jC_k)$, i.e. the probability that a read would have a mutation at position $j$ given that it contained position $j$ and came from cluster $k$; and the observable mutation rates ($m_{jk}$) are $P(M_j|S_jC_kG_g)$, i.e. the probability that a read would have a mutation at position $j$ given that it contained position $j$, came from cluster $k$, and had no two mutations closer than $g$ bases. Using these definitions and Bayes' theorem yields a probabilistic formula for $m_{jk}$:

$$m_{jk} = P(M_j|S_jC_kG_g) = P(M_j|S_jC_k)\frac{P(G_g|S_jM_jC_k)}{P(G_g|S_jC_k)} = \mu_{jk}\frac{P(G_g|S_jM_jC_k)}{P(G_g|S_jC_k)}$$

The term $P(G_g|S_jC_k)$ is the probability that a read would have no two mutations closer than $g$ bases given that it contained position $j$ and came from cluster $k$. It can be computed using $P(G_g|E_{ab}C_k)$ (abbreviated $d_{abk}$): the probability that a read would contain no two mutations closer than $g$ bases given that its 5' and 3' coordinates are $a$ and $b$, repectively ($1 \leq a \leq b \leq L$), and that it came from cluster $k$. If position $b$ were mutated (probability $\mu_{bk}$), then the read would contain no two mutations closer than $g$ bases if and only if none of the $g$ bases preceding $b$ (i.e. positions $b-g$ to $b-1$, inclusive) were mutated (probability $\prod_{j'=\max(b-g,a)}^{b-1}(1-\mu_{j'k})$, abbreviated $w_{\max(b-g,a),b-1,k}$) and two no mutations between positions $a$ and $b-(g+1)$, inclusive, were too close (probability $d_{a,\max(b-(g+1),a),k}$)). If position $b$ were not mutated (probability $1-\mu_{bk}$), then the read would contain no two mutations closer than $g$ bases if and only if no mutations between positions $a$ and $b-1$, inclusive, were too close (probability $d_{a,\max(b-1,a),k}$). These two possibilities generate a recurrence relation:

$$d_{abk} = \mu_{bk} w_{\max(b-g,a),b-1,k} d_{a,\max(b-(g+1),a),k} + (1-\mu_{bk}) d_{a,\max(b-1,a),k}$$

The base case is $d_{abk} = 1$ when $a = b$ because such a read would contain one position and thus be guaranteed to have no two mutations too close. Then, $P(G_g|S_j C_k)$ is the average of $d_{abk}$ over every read that contains position $j$, weighted by the proportions $\eta_{ab}$:

$$P(G_g|S_j C_k) = \frac{\sum_{a=1}^{j} \sum_{b=j}^{L} \eta_{ab} d_{abk}}{\sum_{a=1}^{j} \sum_{b=j}^{L} \eta_{ab}}$$

The term $P(G_g|S_j M_j C_k)$ is the probability that a read would have no two mutations too close given that it contained a mutation at position $j$ and came from cluster $k$. It can be computed using $P(G_g|M_j E_{ab} C_k)$ (abbreviated $f_{abjk}$): the probability that a read would contain no two mutations too close given that position $j$ is mutated ($1 \leq a \leq j \leq b \leq L$), that its 5' and 3' coordinates are $a$ and $b$ (respectively), and that it came from cluster $k$. Because position $j$ is mutated, having no two mutations too close requires that none of the $g$ bases on both sides of position $j$ be mutated. The probability that none of the preceding $g$ positions ($j-g$ to $j-1$) is mutated is $w_{\max(j-g,a),j-1,k}$, while that of the following $g$ positions ($j+1$ to $j+g$) is $w_{j+1,\min(j+g,b),k}$. Upstream of the $g$ bases flanking position $j$ (i.e. positions $a$ to

$j-(g+1)$), the probability that no two mutations are too close is $d_{a,\mathsf{max}(j-(g+1),a),k}$; downstream (i.e. positions $j+(g+1)$ to $b$), the probability is $d_{\mathsf{min}(j+(g+1),b),b,k}$. Since mutations in these four sections are independent, the probability that the read contains no two mutations too close is the product:

$$f_{abjk} = d_{a,\mathsf{max}(j-(g+1),a),k} w_{\mathsf{max}(j-g,a),j-1,k} w_{j+1,\mathsf{min}(j+g,b),k} d_{\mathsf{min}(j+(g+1),b),b,k}$$

Then, $P(G_g|S_jM_jC_k)$ is the average of $f_{abjk}$ over every read that contains position $j$, weighted by the proportions $\eta_{ab}$.

$$P(G_g|S_jM_jC_k) = \frac{\sum_{a=1}^{j}\sum_{b=j}^{L}\eta_{ab}f_{abjk}}{\sum_{a=1}^{j}\sum_{b=j}^{L}\eta_{ab}}$$

Combining the above results yields an explicit formula for $m_{jk}$:

$$m_{jk} = \mu_{jk}\frac{\sum_{a=1}^{j}\sum_{b=j}^{L}\eta_{ab}f_{abjk}}{\sum_{a=1}^{j}\sum_{b=j}^{L}\eta_{ab}d_{abk}}$$

#### Deriving end coordinate proportions of reads with no two mutations too close

The total proportions ($\eta_{ab}$) of reads aligned to 5' and 3' coordinates $a$ and $b$, respectively, are $P(E_{ab})$; and the proportions of reads with no two mutations too close that align with coordinates $a$ and $b$ ($e_{abk}$) are $P(E_{ab}|G_gC_k)$. Note that, while reads are assumed to come from the same distribution of coordinates ($\eta_{ab}$) regardless of their cluster $k$, the observable distribution of coordinates ($e_{abk}$) varies by cluster because P($G_gC_k$) depends on $k$. Using these definitions and Bayes' theorem yields a probabilistic formula for $e_{abk}$:

$$e_{abk} = P(E_{ab}|G_gC_k) = P(G_g|E_{ab}C_k)\frac{P(E_{ab}|C_k)}{P(G_g|C_k)} = d_{abk}\frac{\eta_{ab}}{P(G_g|C_k)}$$

The term $P(G_g|C_k)$ is the probability that a read would have no two mutations too close given that it came from cluster $k$. It can be computed as an average of $P(G_g|E_{ab}C_k)$ (i.e. $d_{abk}$) over all coordinates $a$ and $b$ (such that $1 \leq a \leq b \leq L$), weighted by the proportion of each coordinate, $P(E_{ab})$ (i.e. $\eta_{ab}$):

$$P(G_g|C_k) = \frac{\sum_{a=1}^{L}\sum_{b=a}^{L}\eta_{ab}d_{abk}}{\sum_{a=1}^{L}\sum_{b=a}^{L}\eta_{ab}} = \sum_{a=1}^{L}\sum_{b=a}^{L}\eta_{ab}d_{abk}$$

This expression is already normalized because $\sum_{a=1}^{L}\sum_{b=a}^{L}\eta_{ab} = 1$, by definition.

Combining the above results yields an explicit formula for $e_{abk}$:

$$e_{abk} = \frac{\eta_{ab}d_{abk}}{\sum_{a'=1}^{L}\sum_{b'=a'}^{L}\eta_{a'b'}d_{a'b'k}}$$

#### Deriving cluster proportions of reads with no two mutations too close

The proportion of total reads in cluster $k$ is $\pi_k = P(C_k)$. The proportion among only reads with no two mutations closer than $g$ bases is

$$p_k = P(C_k|G_g) = P(G_g|C_k)\frac{P(C_k)}{P(G_g)} = \pi_k\frac{\sum_{a=1}^{L}\sum_{b=a}^{L}\eta_{ab}d_{abk}}{P(G_g)}$$

The term $P(G_g)$ is the probability that a read from any cluster would have no two mutations closer than $g$ bases and can be solved for by leveraging that the cluster proportions ($p_k$) must sum to 1:

$$1 = \sum_{k=1}^{K}p_k = \sum_{k=1}^{K}\pi_k\frac{\sum_{a=1}^{L}\sum_{b=a}^{L}\eta_{ab}d_{abk}}{P(G_g)} = \frac{1}{P(G_g)}\sum_{k=1}^{K}\pi_k\sum_{a=1}^{L}\sum_{b=a}^{L}\eta_{ab}d_{abk}$$

$$P(G_g) = \sum_{k=1}^{K}\pi_k\sum_{a=1}^{L}\sum_{b=a}^{L}\eta_{ab}d_{abk}$$

The result is an explicit formula for $p_k$:

$$p_k = \frac{\pi_k\sum_{a=1}^{L}\sum_{b=a}^{L}\eta_{ab}d_{abk}}{\sum_{k'=1}^{K}\pi_{k'}\sum_{a=1}^{L}\sum_{b=a}^{L}\eta_{ab}d_{abk'}}$$

#### Solving total mutation rates and cluster and coordinate proportions

The observed mutation rates ($m_{jk}$), end coordinate proportions ($e_{abk}$), and cluster proportions ($p_k$) can be calculated as weighted averages over the $N$ reads with no two mutations too close:

$$m_{jk} = \frac{\sum_{i=1}^{N} z_{ik} x_{ij}}{\sum_{i=1}^{N} z_{ik} s_{ij}}$$

$$e_{abk} = \frac{\sum_{i=1}^{N} z_{ik} y_{abi}}{\sum_{i=1}^{N} z_{ik}}$$

$$p_k = \frac{\sum_{i=1}^{N} z_{ik}}{N}$$

where $s_{ij}$ is $1$ if read $i$ contains position $j$, otherwise $0$; $x_{ij}$ is $1$ if read $i$ has a mutation at position $j$, otherwise $0$; $y_{abi}$ is $1$ if read $i$ aligns to coordinates $a$ and $b$, otherwise $0$; and $z_{ik}$ is the probability that read $i$ came from cluster $k$.

The original parameters $\mu_{jk}$, $\eta_{abk}$, and $\pi_k$ can be solved by setting the two formulae each for $m_{jk}$, $e_{abk}$, and $p_k$ equal to each other, creating a system of equations:

$$\mu_{jk} \frac{\sum_{a=1}^{j} \sum_{b=j}^{L} \eta_{ab} f_{abjk}}{\sum_{a=1}^{j} \sum_{b=j}^{L} \eta_{ab} d_{abk}} = m_{jk} = \frac{\sum_{i=1}^{N} z_{ik} x_{ij}}{\sum_{i=1}^{N} z_{ik} s_{ij}}$$

$$\eta_{ab} \frac{d_{abk}}{\sum_{a'=1}^{L} \sum_{b'=a'}^{L} \eta_{a'b'} d_{a'b'k}} = e_{ab} = \frac{\sum_{i=1}^{N} z_{ik} y_{abi}}{\sum_{i=1}^{N} z_{ik}}$$

$$\pi_k \frac{\sum_{a=1}^{L} \sum_{b=a}^{L} \eta_{ab} d_{abk}}{\sum_{k'=1}^{K} \pi_{k'} \sum_{a=1}^{L} \sum_{b=a}^{L} \eta_{ab} d_{abk'}} = p_k = \frac{\sum_{i=1}^{N} z_{ik}}{N}$$

Solving this entire system at once has proven computationally impractical for all but extremely short sequences. A more feasible approach is to first solve for $\mu_{jk}$ given an initial guess for $\eta_{ab}$, next solve for $\eta_{ab}$ given the updated $\mu_{jk}$, then solve for $\pi_k$ given the updated $\mu_{jk}$ and $\eta_{ab}$, and iterate until all three sets of parameters converge.

Even assuming every $\eta_{ab}$ is a constant, these equations are still too complex to solve for $\mu_{jk}$ analytically because $d_{abk}$ and $f_{abjk}$ also depend on $\mu_{jk}$ (as well as on other $\mu$ variables). Thus, every $\mu_{jk}$ is solved for numerically by rearranging each equation to

$$\mu_{jk}\frac{\sum_{a=1}^{j}\sum_{b=j}^{L}\eta_{ab}f_{abjk}}{\sum_{a=1}^{j}\sum_{b=j}^{L}\eta_{ab}d_{abk}} - m_{jk} = 0$$

and applying the Netwon-Krylov method [85] implemented in SciPy [80].

Once every $\mu_{jk}$ has been solved for, every $\eta_{ab}$ can be updated. Because $d_{abk}$ does not depend on $\eta_{ab}$ (except indirectly through the $\mu_{jk}$ parameters, which are now assumed to be constants), each equation can be rearranged to

$$\eta_{ab} = \frac{e_{ab}}{d_{abk}}\sum_{a'=1}^{L}\sum_{b'=a'}^{L}\eta_{a'b'}d_{a'b'k}$$

Leveraging that $\sum_{a=1}^{L}\sum_{b=a}^{L}\eta_{ab} = 1$, by definition, leads to

$$\sum_{a=1}^{L}\sum_{b=a}^{L}\frac{e_{ab}}{d_{abk}}\sum_{a'=1}^{L}\sum_{b'=a'}^{L}\eta_{a'b'}d_{a'b'k} = 1$$

$$\sum_{a'=1}^{L}\sum_{b'=a'}^{L}\eta_{a'b'}d_{a'b'k} = \frac{1}{\sum_{a=1}^{L}\sum_{b=a}^{L}\frac{e_{ab}}{d_{abk}}}$$

and finally a closed-form expression for each $\eta_{ab}$ given $\mu_{jk}$ (and hence $d_{abk}$) and $e_{abk}$:

$$\eta_{ab} = \frac{\frac{e_{ab}}{d_{abk}}}{\sum_{a'=1}^{L}\sum_{b'=a'}^{L}\frac{e_{a'b'}}{d_{a'b'k}}}$$

This equation should theoretically yield the same value of $\eta_{ab}$ for every $k$. In practice, the values will differ due to inexactness in floating-point arithmetic. Thus, the consensus value of $\eta_{ab}$ is taken to be the average $\eta_{ab}$ over every $k$, weighted by $\pi_k$:

$$\eta_{ab} = \sum_{k=1}^{K}\pi_k\frac{\frac{e_{ab}}{d_{abk}}}{\sum_{a'=1}^{L}\sum_{b'=a'}^{L}\frac{e_{a'b'}}{d_{a'b'k}}}$$

With updated values of $\mu_{jk}$ and $\eta_{ab}$, $\pi_k$ can also be solved. The above equations can be rearranged to

$$\pi_k = p_k\frac{\sum_{k'=1}^{K}\pi_{k'}\sum_{a=1}^{L}\sum_{b=a}^{L}\eta_{ab}d_{abk'}}{\sum_{a=1}^{L}\sum_{b=a}^{L}\eta_{ab}d_{abk}}$$

Given that $\sum_{k=1}^{K} \pi_k = 1$, by definition:

$$\sum_{k=1}^{K} p_k \frac{\sum_{k'=1}^{K} \pi_{k'} \sum_{a=1}^{L} \sum_{b=a}^{L} \eta_{ab} d_{abk'}}{\sum_{a=1}^{L} \sum_{b=a}^{L} \eta_{ab} d_{abk}} = 1$$

$$\sum_{k'=1}^{K} \pi_{k'} \sum_{a=1}^{L} \sum_{b=a}^{L} \eta_{ab} d_{abk'} = \frac{1}{\sum_{k=1}^{K} \frac{p_k}{\sum_{a=1}^{L} \sum_{b=a}^{L} \eta_{ab} d_{abk}}}$$

which leads to a closed-form expression for each $\pi_k$ given $\mu_{jk}$ (and hence $d_{abk}$), $\eta_{ab}$, and $p_k$:

$$\pi_k = \frac{\frac{p_k}{\sum_{a=1}^{L} \sum_{b=a}^{L} \eta_{ab} d_{abk}}}{\sum_{k'=1}^{K} \frac{p_{k'}}{\sum_{a=1}^{L} \sum_{b=a}^{L} \eta_{ab} d_{abk'}}}$$

### Clustering reads with the expectation-maximization algorithm

Let $N$ reads from $K$ clusters align to a reference sequence of length $L$. Let the proportion of reads whose 5' and 3' ends align, respectively, to coordinates $a$ and $b$ ($1 \leq a \leq b \leq L$) be $\eta_{ab}$ (assuming these proportions are equal for all clusters). Let the mutation rate of base $j$ ($1 \leq j \leq L$) in cluster $k$ ($1 \leq k \leq K$) be $\mu_{jk}$. Let the proportion of cluster $k$ in the ensemble be $\pi_k$.

#### Maximization step

The maximization step updates the parameters ($\mu_{jk}$, $\eta_{ab}$, and $\pi_k$) using the current cluster memberships ($z_{ik}$). The observed estimates of the parameters $m_{jk}$, $e_{ab}$, and $p_k$ are first computed; then, the underlying parameters $\mu_{jk}$, $\eta_{ab}$, and $\pi_k$ are solved for as described in 10.1.4.

#### Expectation step

The expectation step updates the cluster memberships ($z_{ik}$) and the likelihood function ($L$) using the current parameters ($\mu_{jk}$, $\eta_{ab}$, and $\pi_k$). Each cluster membership is defined as the probability that read $i$ came from cluster $k$ given its

5'/3' end coordinates ($E_{ab}$) and mutations ($M$) and given that no two mutations are too close ($G_g$): $z_{ik} = P(C_k|E_{ab}MG_g)$. The likelihood of the model ($L$) is the product of the marginal probability ($L_i$) of observing each read $i$ from any cluster: $L_i = P(E_{ab}M|G_g)$. Both $L_i$ and $z_{ik}$ can be expressed in terms of the joint probability ($L_{ik} = P(E_{ab}MC_k|G_g)$) of observing each read $i$ from each cluster $k$:

$$L_i = P(E_{ab}M|G_g) = \sum_{k=1}^{K} P(E_{ab}MC_k|G_g) = \sum_{k=1}^{K} L_{ik}$$

$$z_{ik} = P(C_k|E_{ab}MG_g) = \frac{P(E_{ab}MC_kG_g)}{P(E_{ab}MG_g)} = \frac{P(E_{ab}MC_k|G_g)}{P(E_{ab}M|G_g)} = \frac{L_{ik}}{L_i}$$

To derive a formula for $L_{ik}$, it can be factored into three parts using the chain rule for probability:

$$L_{ik} = P(E_{ab}MC_k|G_g) = \frac{P(E_{ab}MC_kG_g)}{P(G_g)} = P(M|E_{ab}C_kG_g)P(E_{ab}|C_kG_g)P(C_k|G_g)$$

The first part – the probability that a read would have the specific mutations $x_{ij}$ given that its 5'/3' end coordinates are $a$ and $b$ (respectively), it comes from cluster $k$, and no two mutations are too close – is the product over every position $j$ from $a$ to $b$ of the probability of a mutation ($\mu_{jk}$) if read $i$ is mutated at position $j$ ($x_{ij} = 1$), otherwise ($x_{ij} = 0$) the probability of no mutation ($1 - \mu_{jk}$), normalized by the probability that no two mutations would be too close ($d_{abk}$):

$$P(M|E_{ab}C_kG_g) = \frac{1}{d_{abk}} \prod_{j=a}^{b} \mu_{jk}^{x_{ij}} (1 - \mu_{jk})^{(1-x_{ij})}$$

The second part, $P(E_{ab}|C_kG_g) = e_{abk}$, can be calculated from the parameters $\mu_{jk}$, $\eta_{ab}$, and $\pi_k$, as explained in 10.1.2. Likewise, the third part, $P(C_k|G_g) = p_k$, can also be calculated from the parameters, as explained in 10.1.3. Combining all parts yields a formula for $L_{ik}$ in terms of the parameters $\mu_{jk}$, $\eta_{ab}$, and $\pi_k$ and of their derived values $d_{abk}$, $e_{abk}$, and $p_k$:

$$L_{ik} = p_k \frac{e_{abk}}{d_{abk}} \prod_{j=a}^{b} \mu_{jk}^{x_{ij}} (1 - \mu_{jk})^{(1-x_{ij})}$$

The formula for the total likelihood of the model and its parameters follows:

$$L(\mu, \eta, \pi) = \prod_{i=1}^{N} L_i = \prod_{i=1}^{N} \sum_{k=1}^{K} p_k \frac{e_{abk}}{d_{abk}} \prod_{j=a}^{b} \mu_{jk}^{x_{ij}} (1 - \mu_{jk})^{(1-x_{ij})}$$

### Supplementary Tables

Table 1: **Sequences of the antisense oligonucleotides (ASOs) targeting the 2,924 nt segment of SARS-CoV-2 RNA.**

| Group | ASO | Sequence |
|---|---|---|
| 1 | 1 | GGCAGCACAAGACATCTGTCGTAGTGCAACAGGACTAA- GCTCATTATT |
| | 2 | TGTAGTAAGCTAACGCATTGTCATCAGTGCAAGCAGTTT- GTGTAGTACC |
| | 3 | TGTAAATCGGATAACAGTGCAAGTACAAACCTACCTCCC- TTTGTTGTGT |
| | 4 | GATAGTACCAGTTCCATCACTCTTAGGGAATCTAGCCCA- TTTCAAATCC |
| | 5 | CTTTAGGTGTGTCTGTAACAAACCTACAAGGTGGTTCCA- GTTCTGTATA |
| 2 | 1 | ATACCTCTATTTAGGTTGTTTAATCCTTTAATAAAGTATAA- ATACTTCACTTTAGGAC |
| | 2 | CACTTCTGTTGCATTACCAGCTTGTAGACGTACTGTGGC- AGCTAAACTACCAAGTACC |
| | 3 | AAGCTTTAGCAGCATCTACAGCAAAAGCACAGAAAGATA- ATACAGTTGAATTGGCAGG |
| | 4 | CACAACATCTTAACACAATTAGTGATTGGTTGTCCCCCA- CTAGCTAGATAATCTTTGT |
| 3 | 1 | GATCCATATTGGCTTCCGGTGTAACTGTTATTGCCTGAC- CAGTACCAGTGTGTGTA |
| | 2 | ATGATCTATGTGGCAACGGCAGTACAGACAACACGATG- CACCACCAAAGGATTCTT |

Continued on next page

Table 1: **Sequences of the antisense oligonucleotides (ASOs) targeting the 2,924 nt segment of SARS-CoV-2 RNA.** (Continued)

| | | |
|---|---|---|
| | 3 | GTTGTAGGTATTTGTACATACTTACCTTTTAAGTCACAAA-ATCCTTTAGGATTTGG |
| | 4 | CCGCAGACGGTACAGACTGTGTTTTTAAGTGTAAAACCC-ACAGGGTCATTAGCACAA |
| 4 | 1 | CTGAAGCATGGGTTCGCGGAGTTGATCACAACTACAGC-CATAACCTTTCCACATA |
| | 2 | GGAAGCGACAACAATTAGTTTTTAGGAATTTAGCAAAAC-CAGCTACTTTATCATTG |
| 5 | 1 | TGTCTCTTAACTACAAAGTAAGAATCAATTAAATTGTCAT-CTTCGTCCTTTTCTT |
| | 2 | GACAATCCTTAAGTAAATTATAAATTGTTTCTTCATGTTG-GTAGTTAGAGAAAGTG |
| | 3 | GGTACCATGTCACCGTCTATTCTAAACTTAAAGAAGTCA-TGTTTAGCAACAGCTG |
| | 4 | AAGCATAGACGAGGTCTGCCATTGTGTATTTAGTAAGAC-GTTGACGTGATATATGT |
| 6 | 1 | TGTATGTGACAAGTATTTCTTTTAATGTGTCACAATTACC-TTCATCAAAATGCCTTA |
| | 2 | GGTTTTCTACAAAATCATACCAGTCCTTTTTATTGAAATA-ATCATCATCACAACAAT |
| | 3 | TTAACAAAGCTTGGCGTACACGTTCACCTAAGTTGGCGT-ATACGCGTAATATATCTG |
| | 4 | ATGTCAGTACACCAACAATACCAGCATTTCGCATGGCAT-CACAGAATTGTACTGTTT |

Continued on next page

Table 1: **Sequences of the antisense oligonucleotides (ASOs) targeting the 2,924 nt segment of SARS-CoV-2 RNA.** (Continued)

| | | |
|---|---|---|
| 7 | 1 | GTTTGTATGAAATCACCGAAATCATACCAGTTACCATTG-AGATCTTGATTATCTA |
| | 2 | TAGGCATTAACAATGAATAATAAGAATCTACAACAGGAA-CTCCACTACCTGGCGTG |
| | 3 | GTTAAGTCAGTGTCAACATGTGACTCTGCAGTTAAAGCC-CTGGTCAAGGTTAATA |
| | 4 | TTAACCTCTCTTCCGTGAAGTCATATTTTAACAAATCCCA-CTTAATGTAAGGCTTT |
| 8 | 1 | AACACAATTTGGGTGGTATGTCTGATCCCAATATTTAAAA-TAACGGTCAAAGAGTT |
| | 2 | GAGAATAAAACATTAAAGTTTGCACAATGCAGAATGCAT-CTGTCATCCAAACAGTT |
| | 3 | CATCAACAAATATTTTTCTCACTAGTGGTCCAAAACTTGT-AGGTGGGAACACTGTA |
| | 4 | ATGTACAACACCTAGCTCTCTGAAGTGGTATCCAGTTGA-AACTACAAATGGAACAC |
| 9 | 1 | TACACAAGTAATTCCTTAAAACTAAGTCTAGAGCTATGTA-AGTTTACATCCTGATT |
| | 2 | TGCGTTTATCTAGTAATAGATTACCAGAAGCAGCGTGCA-TAGCAGGGTCAGCAGCA |
| | 3 | TTTGACAGTTTGAAAAGCAACATTGTTAGTAAGTGCAGC-TACTGAAAAGCACGTAG |
| | 4 | CTTAAAGAAACCCTTAGACACAGCAAAGTCATAGAAGTC-TTTGTTAAAATTACCGGG |

Continued on next page

Table 1: **Sequences of the antisense oligonucleotides (ASOs) targeting the 2,924 nt segment of SARS-CoV-2 RNA.** (Continued)

| | | |
|---|---|---|
| 10 | 1 | CAGCATTACCATCCTGAGCAAAGAAGAAGTGTTTTAATT-CAACAGAACTTCCTTC |
| | 2 | CTGATATCACACATTGTTGGTAGATTATAACGATAGTAGT-CATAATCGCTGATAG |
| | 3 | ACCATCGTAACAATCAAAGTACTTATCAACAACTTCAACT-ACAAATAGTAGTTGT |
| | 4 | AACCAGCTGATTTGTCTAGGTTGTTGACGATGACTTGGT-TAGCATTAATACAGCC |
| 11 | 1 | CCTCATAACTCATTGAATCATAATAAAGTCTAGCCTTACC-CCATTTATTAAATGGAA |
| | 2 | ATTTGAGTTATAGTAGGGATGACATTACGTTTTGTATATG-CGAAAAGTGCATCTTGAT |
| | 3 | GAGACACCAGCTACGGTGCGAGCTCTATTCTTTGCACTA-ATGGCATACTTAAGATTC |
| | 4 | GGCTATTGATTTCAATAATTTTTGATGAAACTGTCTATTG-GTCATAGTACTACAGATA |
| 12 | 1 | CAACCACCATAGAATTTGCTTGTTCCAATTACTACAGTA-GCTCCTCTAGTGGC |
| | 2 | CCATAAGGTGAGGGTTTTCTACATCACTATAAACAGTTTT-TAACATGTTGTGC |
| | 3 | CATAATTCTAAGCATGTTAGGCATGGCTCTATCACATTTA-GGATAATCCCAAC |
| | 4 | ACGGTGTGACAAGCTACAACACGTTGTATGTTTGCGAG-CAAGAACAAGTGAGGC |

Continued on next page

Table 1: **Sequences of the antisense oligonucleotides (ASOs) targeting the 2,924 nt segment of SARS-CoV-2 RNA.** (Continued)

| | | |
|---|---|---|
| 13 | 1 | ACACATGACCATTTCACTCAATACTTGAGCACACTCATTA-GCTAATCTATAGAA |
| | 2 | AGTTGTGGCATCTCCTGATGAGGTTCCACCTGGTTTAAC-ATATAGTGAACCGCC |
| | 3 | ATTAACATTGGCCGTGACAGCTTGACAAATGTTAAAAAC-ACTATTAGCATAAGC |
| | 4 | TAAATTGCGGACATACTTATCGGCAATTTTGTTACCATCA-GTAGATAAAAGTGC |

Table 2: **Sequences of the forward (F) and reverse (R) primers for amplifying the target site of each ASO group in the 2,924 nt segment of SARS-CoV-2 RNA.**

| Group | Primer | Sequence |
|---|---|---|
| 1 | F | AATAATGAGCTTAGTCCTGTTGCACTACG |
| | R | AGGTTGTTTAATCCTTTAATAAAGTATAAATACTTCACT-TTAGG |
| 2 | F | ACCTTGTAGGTTTGTTACAGACACACCTAA |
| | R | TTGCCTGACCAGTACCAGTGTGTG |
| 3 | F | GGACAACCAATCACTAATTGTGTTAAGATGTTG |
| | R | TCACAACTACAGCCATAACCTTTCCACA |
| 4 | F | CTTAAAAACACAGTCTGTACCGTCTGC |
| | R | GTAAGAATCAATTAAATTGTCATCTTCGTCCTTTTC |
| 5 | F | TGCTAAATTCCTAAAAACTAATTGTTGTCGCTT |
| | R | ATGTGTCACAATTACCTTCATCAAAATGCCT |
| 6 | F | CAATGGCAGACCTCGTCTATGC |
| | R | GAAATCATACCAGTTACCATTGAGATCTTGATTATC |
| 7 | F | CGAAATGCTGGTATTGTTGGTGTACTGAC |
| | R | GTCTGATCCCAATATTTAAAATAACGGTCAAAGAG |
| 8 | F | TGTTAAAATATGACTTCACGGAAGAGAGGTT |
| | R | AAGTCTAGAGCTATGTAAGTTTACATCCTGA |
| 9 | F | CCACTTCAGAGAGCTAGGTGTTGTAC |
| | R | CAAAGAAGAAGTGTTTTAATTCAACAGAACTTCCT |
| 10 | F | TGACTTTGCTGTGTCTAAGGGTTTCTTTA |
| | R | CATAATAAAGTCTAGCCTTACCCCATTTATTAAATGG |
| 11 | F | CGTCAACAACCTAGACAAATCAGCTGG |

Continued on next page

Table 2: **Sequences of the forward (F) and reverse (R) primers for amplifying the target site of each ASO group in the 2,924 nt segment of SARS-CoV-2 RNA.** (Continued)

| | | |
|---|---|---|
| | R | TTCCAATTACTACAGTAGCTCCTCTAGTG |
| 12 | F | GACCAATAGACAGTTTCATCAAAAATTATTGAAATCAA-TAG |
| | R | ATACTTGAGCACACTCATTAGCTAATCTATAG |
| 13 | F | ACAACGTGTTGTAGCTTGTCACACC |
| | R | TAAATTGCGGACATACTTATCGGCAATTTTG |

Table 3: **Sequences of the antisense oligonucleotides (ASOs) targeting the 1,799 nt segment of SARS-CoV-2 genomic RNA.** A plus sign (+) indicates that the following nucleotide is locked nucleic acid (LNA).

| ASO | Sequence |
|---|---|
| Anti-LS1 | GTAATTC+CTTAAAA+CTAAG |
| Anti-LS2a | TGAAA+AGCAA+CATTGTT |
| Anti-LS2b | TA+CCGGGTTTGACAG |
| Anti-LS3b | A+CCCTTAGACACAGCA |
| Anti-AS1 | TGGGTTCGCG+GAGTTG |
| Anti-PS2-overlap | GT+TAAAATTA+CCG+GG |

Table 4: **PCR primer annealing temperatures for coronavirus gene fragments.**

| **Coronavirus** | **Annealing Temperature (°C)** |
|---|---|
| Bat Coronavirus 1A | 55 |
| Bat Coronavirus BM48-31 | 60 |
| Common Moorhen Coronavirus | 55 |
| Human Coronavirus OC43 | 55 |
| Infectious Bronchitis Virus | 60 |
| MERS Coronavirus | 60 |
| Murine Hepaitis Virus | 60 |
| SARS Coronavirus 1 | 60 |
| SARS Coronavirus 2 | 55 |
| Transmissible Gastroenteritis Virus | 55 |

Table 5: **Sequences of the forward (F), forward with T7 promoter (F+T7), and reverse (R) primers for amplifying the 239 nt segment of each 1,799 nt segment of coronaviral RNAs.**

| Coronavirus | Primer | Sequence |
|---|---|---|
| Bat Coronavirus 1A | F | GGACCCTATACGGTTTTGT-CTTGAAAA |
| | F+T7 | TAATACGACTCACTATAGGA-CCCTATACGGTTTTGTCTTG-AAAA |
| | R | TTTTACAATAAAGAAAGCAT-CATGCTT |
| Bat Coronavirus BM48-31 | F | GGGTTTTATTCTTAGAAACA-CAGTCTG |
| | F+T7 | TAATACGACTCACTATAGG-GTTTTATTCTTAGAAACACA-GTCTG |
| | R | GGAGTCTAATAAGTTGCCC-TCTTCATC |
| Common Moorhen Coronavirus | F | GGATAAAGATAAGGAACCT-GTTTCTTT |
| | F+T7 | TAATACGACTCACTATAGGA-TAAAGATAAGGAACCTGTTT-CTTT |
| | R | ACTATTAGGTATTGGCAAAT-TAATGCG |
| Human Coronavirus OC43 | F | GGCTGTGTCTTATGTTTTGA-CACATGA |

Continued on next page

Table 5: **Sequences of the forward (F), forward with T7 promoter (F+T7), and reverse (R) primers for amplifying the 239 nt segment of each 1,799 nt segment of coronaviral RNAs.** (Continued)

| | | |
|---|---|---|
| | F+T7 | TAATACGACTCACTATAGG-CTGTGTCTTATGTTTTGACA-CATGA |
| | R | ATCTAATTTATCACCGTTCT-CATCAAC |
| Infectious Bronchitis Virus | F | GGTTTGCACTGTTTGCCAG-TGTTGGAT |
| | F+T7 | TAATACGACTCACTATAGGT-TTGCACTGTTTGCCAGTGTT-GGAT |
| | R | CTCAAGATTTCCATCTTCAG-TATCGCG |
| MERS Coronavirus | F | GGGATTTTGTTTGTCAAATA-CCCCCTG |
| | F+T7 | TAATACGACTCACTATAGG-GATTTTGTTTGTCAAATACC-CCCTG |
| | R | ATGATGCCCTTGGTCATCT-AATTCTAC |
| Murine Hepatitis Virus | F | GGCTGTGTCATATGTGTTG-ACGCATGA |
| | F+T7 | TAATACGACTCACTATAGG-CTGTGTCATATGTGTTGAC-GCATGA |

Continued on next page

Table 5: **Sequences of the forward (F), forward with T7 promoter (F+T7), and reverse (R) primers for amplifying the 239 nt segment of each 1,799 nt segment of coronaviral RNAs.** (Continued)

| | | |
|---|---|---|
| | R | ATCCAACTTGTTGCCGTCC-TCATCTAC |
| SARS Coronavirus 1 | F | GGGTTTTACACTTAGAAACA-CAGTCTG |
| | F+T7 | TAATACGACTCACTATAGG-GTTTTACACTTAGAAACACA-GTCTG |
| | R | AGAGTCTAATAAATTGCCTT-CCTCATC |
| SARS Coronavirus 2 | F | GGGTTTTACACTTAAAAACA-CAGTCTG |
| | F+T7 | TAATACGACTCACTATAGG-GTTTTACACTTAAAAACACA-GTCTG |
| | R | AGAATCAATTAAATTGTCAT-CTTCGTC |
| Transmissible Gastroenteritis Virus | F | GGCAATTCGGTTCTGTATT-GAAAATGA |
| | F+T7 | TAATACGACTCACTATAGG-CAATTCGGTTCTGTATTGAA-AATGA |
| | R | TTTGACAATGTAGTAGGCAT-CATGTTT |

Table 6: **Sequences of the antisense oligonucleotides (ASOs) targeting the 1,799 nt segments of coronaviral RNAs.**

| Coronavirus | ASO | Sequence |
|---|---|---|
| Bat Coronavirus 1A | 1 | CAGGGCTCTAGTCGAGCTGC-ACTAGAGCCCCTTGCTCGTTT-AAATAAGCCTGATCAACAG |
| | 2 | GCAACTTCTTTATTGTAAATAT-CAAAGGCGCGTACAACATGC-TCCGGTTCAGTACCATTA |
| Bat Coronavirus BM48-31 | 1 | GACATCAGTGCTTGTGCCTGT-GCCGCACGGTGTAAGACGGG-CCGCACTTACACCGCAAAC |
| | 2 | TTTTAGGAACTTTGCAAAACC-AGCAACTTTCTCATTATAAATA-TCAAAAGCCCTGTAAAC |
| | 3 | AAAATAGGAGTCTAATAAGTT-GCCCTCTTCATCAACTTCCTG-GAAACGGCAACAATTTGT |
| Common Moorhen Coronavirus | 1 | TGGGGTTCTAGACGGGCATC-ACTAGAACCCTTTACTCGTTT-AAATAAGCTGTATTTTGCA |
| | 2 | GTTATATTATTATGTACATGAA-ACGCCCTTTTTACAATATCCG-GCTGAGTGCCAGACTGT |
| Infectious Bronchitis Virus | 1 | ACATCAAAGGCTCGCTTTACA-ACATCAGGATCACATCCACTA-GCAAGGGGTATCAGCCGA |

Continued on next page

Table 6: **Sequences of the antisense oligonucleotides (ASOs) targeting the 1,799 nt segments of coronaviral RNAs.** (Continued)

| | | |
|---|---|---|
| Murine Hepatitis Virus | 1 | AAGCCACTGGCACAGGGTAC-AAGACGGGCATTTACACTTGT-ACCCCGAATCCGTTTAAAA |
| | 2 | CCAATGCCAGCTCGATTAGCA-TTACAAATGTCAAATGCCCTT-AATTGAACATCAGTGTCC |
| | 3 | AACTTGTTGCCGTCCTCATCT-ACACGCTGGAAGCGGCAGCA-ATTCACTTTATAATACAAA |
| SARS Coronavirus 1 | 1 | TTTGCAAAACCAGCAACTTTT-TCGTTGTAAATATCAAAAGCC-CTGTAGACGACATCAGTA |
| | 2 | TCTAATAAATTGCCTTCCTCAT-CCTTCTCCTGGAAGCGACAG-CAATTAGTTTTTAGGAAC |
| SARS Coronavirus 2 | 1 | GACATCAGTACTAGTGCCTGT-GCCGCACGGTGTAAGACGGG-CTGCACTTACACCGCAAAC |
| | 2 | TTTTAGGAATTTAGCAAAACC-AGCTACTTTATCATTGTAGAT-GTCAAAAGCCCTGTATAC |
| Transmissible Gastroenteritis Virus | 1 | TAAATAACTTTGATCAACAGT-AAAACTCTGCATAGAAGTACG-ATCGCACATGCAACCATT |

Continued on next page

Table 6: **Sequences of the antisense oligonucleotides (ASOs) targeting the 1,799 nt segments of coronaviral RNAs.** (Continued)

| | | |
|---|---|---|
| | 2 | GGTCTGGATCAGTACCATTGC-AGGGTTCTAGTCGAGCTGCA-CTAGAACCCCGCACTCGTT |
